# Region- and variance-based DNA methylation analyses reveal novel disease genes and pathways for systemic lupus erythematosus

**DOI:** 10.1101/2022.11.23.516835

**Authors:** Mengbiao Guo, Ting-You Wang, Jiangshan Jane Shen, Yong-Fei Wang, Yu-Lung Lau, Wanling Yang

**Affiliations:** Department of Paediatrics and Adolescent Medicine, Li Ka Shing Faculty of Medicine, The University of Hong Kong

**Keywords:** systemic lupus erythematosus, methylation variability, T cell receptor, B cell receptor, enhancer, machine learning, disease diagnosis

## Abstract

**Background:** Systemic lupus erythematosus (SLE) is a prototype autoimmune disease with unclear pathogenesis. DNA methylation is an important regulatory mechanism on gene expression, providing a key angle to understand disease mechanisms. To understand the pathways involved in SLE, and to develop biomarkers for its diagnosis and treatment, we analyzed DNA methylation profiles on blood cells from SLE patients and healthy controls.

**Results:** We identified most differentially methylated regions (DMRs) in T cells, while majority of differentially variable sites (DVSs) were found in B cells, featuring hypervariability in enhancers. We observed a prominent T cell receptor (TCR) signaling cluster with consistent hypermethylation and a B cell receptor (BCR) cluster with highly increased variability in SLE. Genes involved in innate immunity were often found hypomethylated, while adaptive immunity genes were featured with hypermethylation. Using a machine learning approach, we identified 60 genes that accurately distinguished SLE patients from healthy individuals, which also showed correlation with disease activities.

**Conclusions:** This study highlights the role of lymphocyte receptor aberrations in the disease and identified a list of genes showing great potential as biomarkers and shedding new light on disease mechanisms, through novel analyses of methylation data.

## Background

Systemic lupus erythematosus (SLE) [1] is a prototypic autoimmune disease with autoantibody production and multi-organ damage. Although recent progress in genome-wide association studies (GWAS) has advanced our understanding of the genetic architecture of this disease, the explained genetic heritability by the confirmed susceptibility variants is only about 30% [2]. On the other hand, studies revealed indispensable roles of epigenetic changes in SLE pathogenesis, especially DNA methylation [3, 4], an epigenetic modification on DNA that has been widely studied in various human diseases [5, 6]. Due to technical limitations, however, most previous epigenetic studies focused only on a single gene or a handful of candidate genes [7], such as *ITGAL, IL-4*, and *CD40L*. In the past, several genome-wide studies of DNA methylation have been performed on SLE as well [8-16]. Absher et al. and Coit et al. reported prominent hypomethylation of genes involved in interferon signaling in blood cells [9, 10]. Imgenberg-Kreuz et al. identified close links between genetic variants and changes in DNA methylation in SLE patients [16].

Although genome-wide epigenetic studies have found thousands of differentially methylated genes, most were based on single-CpG differential analysis (DMC) and the main findings were concentrated on hypomethylated genes, especially those in the type I interferon signaling pathway. We hypothesize that promising insights for SLE pathogenesis could emerge by adopting a recently published algorithm analyzing differentially methylated regions (DMR), which is much more powerful than DMC analysis with comparable false positive rate [17], as well as an algorithm studying DNA methylation variability (DVS) [18].

Here, we present findings from an in-depth analysis of abnormal methylation in SLE patients based on DMR and DVS analyses of published DNA methylation datasets on T cells, B cells, monocytes, and PBMCs. We found hypermethylated genes were significantly overrepresented in those involved in adaptive immunity, especially the T cell receptor (TCR) co-signaling pathways, and genes with higher methylation viability were enriched in the B cell receptor (BCR) signaling pathways. Further, we found DVSs were specifically enriched in enhancers, supported by the enrichment of a list of enhancer-binding transcription factors (TFs) in genes with DVSs, including EP300, CEBPB, KDM1A, CHD7, and PAX5. Finally, using a machine learning approach, we selected a list of CpG sites that accurately predicted SLE disease status and correlated with disease activities.

## Methods

### Data sources

All methylation datasets were downloaded from Gene Expression Omnibus (GEO), including GSE59250 for CD4+ T cells, CD19+ B cells and CD14+ monocytes from SLE patients and controls of European descent [9] and GSE82218 for PBMC from Chinese SLE patients and controls [11]. Rheumatoid arthritis (RA) methylation samples were from GSE71841 for CD4+ T cells [19] and GSE87095 for CD19+ B cells [20]. All the above methylation datasets were generated by HumanMethylation450 BeadChip.

### DMR identification

All methylation signals were preprocessed according to the recommended workflow from Bioconductor R package *wateRmelon*. Probes with Illumina BeadChip signal detection P-value>0.05 were set as missing. Probes with missing data among >1% samples and samples with missing data among >1% probes were removed. Probes on sex chromosomes X were also excluded from further analysis. Then, the DASEN method was adopted to normalize the methylation signals as recommended [21]. Same as previous studies [22], methylation Beta-value were calculated as

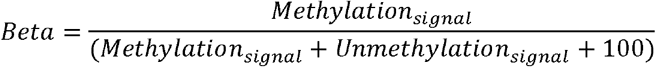

which was converted to M-value by

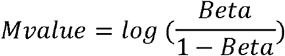

Finally, DMRcate [23] was applied to M-values to identify DMRs of genome-wide significance by a P-value threshold of 1e-7 derived through Bonferroni correction (∼0.05/485,577). Batch, sex, age, and ethnicity were considered as covariates. In addition, with or without removing cross-reactive probes or probes with SNP variants near the CpG sites before calling DMRs had negligible effect on the results.

### DVS identification

Methylation sites with differential variability (DVSs) in CD4+ T cells, CD19+ B cells, and CD14+ monocytes were identified by the iEVORA algorithm [18], by using the preprocessed data prepared for DMR analysis. This method detects DVSs with a differential variation FDR <0.001 and differential average methylation with P-value <0.05 (unadjusted P-value, no genome-wide significance required for DVS).

### Network analysis for DMR and DVS genes based on PPI

After mapping DMR genes to network nodes of the STRING database of protein-protein interaction, a subnetwork was extracted by using an algorithm from the R package XGR [24, 25], consisting of genes with the maximum score (defined as a combination of gene P-values from DMR or DVS analysis). This subnetwork contained both as many most significant genes as possible and a few less significant genes serving as linkers in the subnetwork -- in total, 54 genes and 115 interactions for T cell DMRs and 54 genes and 77 interactions for B cell DVSs. Pathway and disease ontology enrichment were performed with the genes of this subnetwork by using databases of MSigDB.

### TF enrichment in DMRs and DVSs

To analyze the enrichment of TFs bound to the differentially methylated genes (DMGs), 1,183 preprocessed TF binding datasets (TFBS) obtained from chromatin immunoprecipitation experiments (ChIP-Seq) were downloaded from the ENCODE Project website (encodeproject.org). Each TFBS dataset was examined for overlap with DMG promoter-proximal regions (2Kb up- and downstream of TSS) separately for each DMG type (hyper- or hypomethylation) and each cell type. It provided the observed total occurrence (BSdmg) of TFBS among DMGs, with multiple binding sites within the same gene recorded cumulatively. Next, 1,000 simulations by permutation were performed to obtain a background distribution of TFBS occurrence among DMGs under random chance (BSrandom), and subsequently fold-enrichment (BSdmg divided by average of BSrandom values) and empirical P-value (proportion of BSrandom values that are larger than BSdmg) were calculated. Each simulation process starts by randomly sampling the same number of genes as each set of DMGs from human protein-coding RefSeq genes, and the BSrandom value was recorded as the total TFBS occurrence for these sampled genes.

### Machine learning on DMR genes to distinguish SLE patient samples from controls

Machine learning was used to distinguish SLE patients from healthy individuals based on methylation data. First, LASSO (R package glmnet) was applied to select a group of independently contributing CpGs within DMRs, by using methylation M-values of all CpGs within DMRs as input. It achieved this by training a generalized linear model and performing feature selection by L1-norm penalized regression and cross-validation. The performance of the model was then tested by its application to SLE datasets from various cell types, using methylation signals from the same set of selected CpGs in the corresponding samples. Before being fed into LASSO, methylation M-values were standardized to have mean of zero and variance of one, for both training and testing datasets. M-score for each sample was obtained by a weighted sum of standardized methylation of each selected CpG multiplied by model coefficients.

## Results

### Identification of genome-wide differential methylation

DNA methylation of CpG sites has been shown to be strongly correlated with each other in a window of as large as 1,000 base pairs [26], and Jaffe et al. has demonstrated that, compared to the single-CpG analysis, large improvement on true signal detection can be achieved by integrating this information into statistical models [17]. By analyzing a publicly available dataset of DNA methylation for SLE [9], we identified differentially methylated regions (DMRs) across the genome in blood cells from SLE patients of European descent (**Table 1**), and validated the findings using additional datasets [11].

**Table 1.** Summary of samples used and DMRs identified in different cell types.

|  | Ethnicity | Sample |  | Promoter (%)<sup>1</sup> |  | Non-Promoter (%) |  |
| --- | --- | --- | --- | --- | --- | --- | --- |
|  |  | S | Contr | Hypo<sup>2</sup> | Hyper<sup>3</sup> | Hypo | Hyper |
|  |  | L | ol |  |  |  |  |
|  |  | E |  |  |  |  |  |
| **T cell (CD4)** [9] | European | 54 | 56 | 1270 (34) | 600 (16) | 1060 (28) | 790 (22) |
| **B cell (CD19)** [9] | European | 48 | 56 | 114 (31) | 103 (28) | 71 (19) | 83 (22) |
| **Monocyte (CD14)** [9] | European | 27 | 28 | 52 (48) | 18 (16) | 15 (14) | 24 (22) |
| **PBMC** [11] | Chinese | 15 | 25 | 2570 (36) | 1334 (18) | 1551 (21) | 1775 (25) |
| **Naïve T cell (CD4)** [10] | European | 74 | - | - | - | - | - |

<sup>1</sup> Promoter region: defined as 2000bp upstream and downstream of transcription start site (TSS);
<sup>2</sup> Hypo: hypo-methylated; <sup>3</sup> Hyper: hyper-methylated;
<sup>4</sup> used for correlation between disease activity and methylation;

Briefly, with a P-value threshold of 1.0e-7 (genome-wide significance by Bonferroni correction), we found 3,720 DMRs in T cells (CD4+), of which about two-thirds (62%) were hypomethylated, consistent with previous reports [15]. Among these DMRs, 50% (1,870) were in the promoter-proximal region of genes (defined as a 4-kb region centered on the transcription start site (TSS) for each gene), while others fell into gene bodies or intergenic regions. We found fewer DMRs passed genome-wide significance threshold in B cells (CD19+) and monocytes (CD14+), namely 371 and 109 for these two cell types, respectively. This was unlikely to be only due to detection power, as at least for B cells, similar number of samples was available for analysis (**Table 1**). Half of DMRs found in B lymphocytes and monocytes were also identified in T cells, as shown in **Fig. 1a** (see also Additional file 1: **Tables S1-3**), indicating highly shared methylation aberrations among these cell types. Moreover, a large fraction of these DMRs were validated in peripheral blood mononuclear cells (PBMC, Additional file 1: **Table S4**, Additional file 2: **Supplementary Results**). These DMRs were further compared to the significant CpGs reported in the original publication on this dataset (Additional file 2: **Supplementary Results**) [9]. More than three fourths of the DMRs were novel from this study, largely detected from T cells (**Fig. 1b**), which was expected given the advantages of region-based statistical models over CpG-based methods [17]. These novel DMRs tend to have slightly lower significance (**Fig. 1c**), but as expected, they generally contain more CpG probes (**Fig. 1c, t-test P-value= 6e-13**), which is the basis of the higher sensitivity of the DMRcate algorithm.

**Fig. 1.**
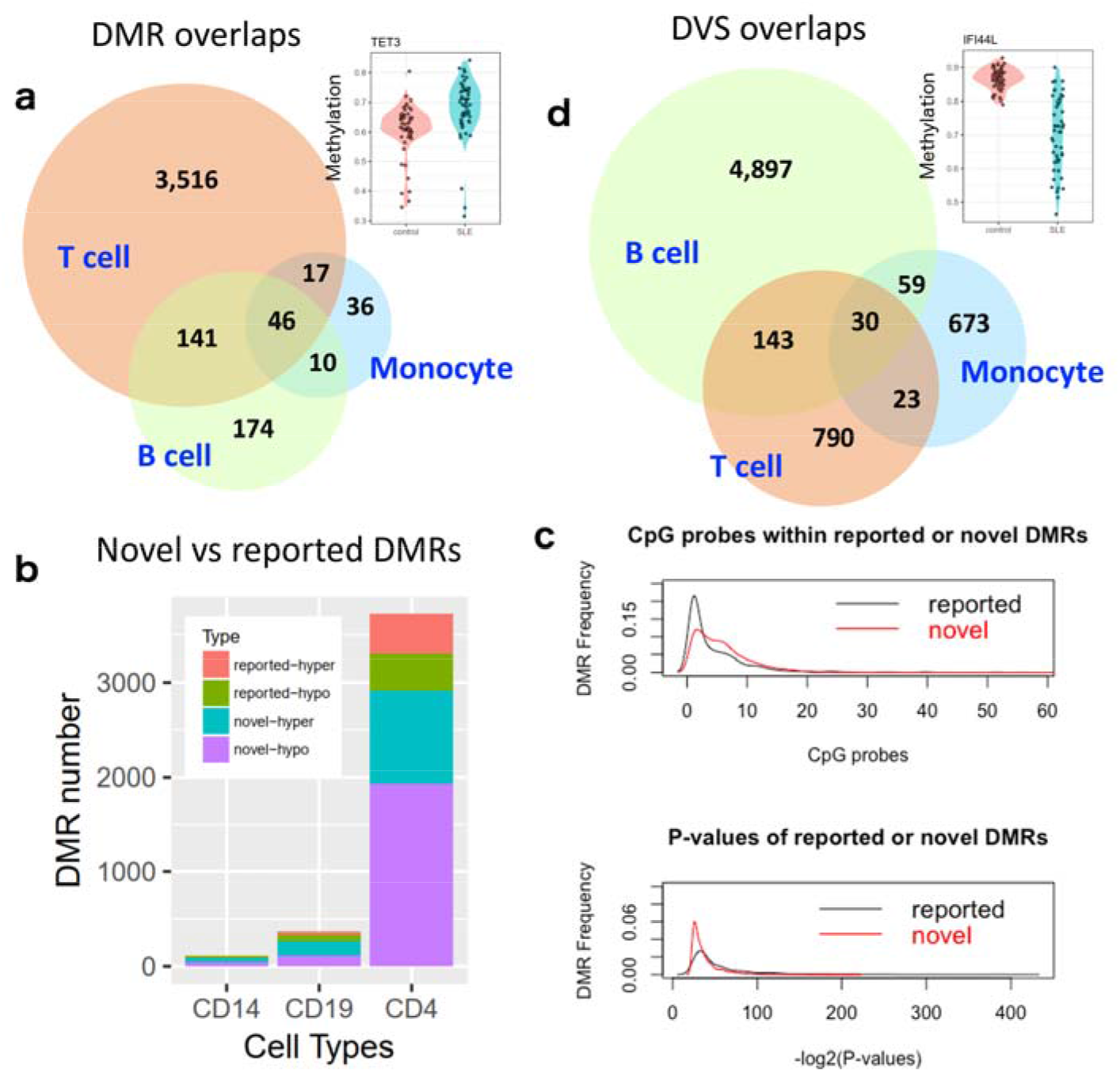
DMR and DVS overlaps between cell types. (**a**) Overlapping of DMRs from T cells, B cells and monocytes. Most DMRs were identified in T cells and half of DMRs in B cell and monocytes were also found in T cells. A typical example of differentially methylated CpG in a DMR of *TET3* was shown (inset). (**b**) Comparison of DMRs with genome-wide significant CpGs reported in the original study on the same dataset. (**c**) Comparison of reported and novel DMRs. Novel DMRs tend to have lower significance, but contain more CpG probes (P-value 6e-13). (**d**) Overlapping of DVSs from T cells, B cells and monocytes. Most DVSs were found in B cells, while only a small number of DVSs from T cells and monocytes were also found in B cells. A typical example of DVS in gene *IFI44L* was shown (inset).

Seeking to further understand the large discrepancy in DMR numbers between T cells and B cells, we performed differential variability analysis using the iEVORA algorithm [18]. Differentially variable sites (DVSs) were defined as CpGs presenting different variance of DNA methylation between patient group and control group, in contrast to the comparison between average levels of methylation (DMR). They have been well studied in both cancer and some complex diseases [18, 27]. Interestingly, we observed far more methylation variable sites in B cells of SLE patients than in T cells (**Fig. 1d**, Additional file 2: **Supplementary Fig. 1**, Additional file 1: **Tables S5-7**). More than half (56%) of the DMRs from B cells harbored at least one DVS (with a median number of 2), compared with only 4.3% in T cells (with a median of 1 DVS). Compared to DMRs, DVSs were far more likely to be cell-type specific. In addition, we found that 26% (44/166) of the B cell DMCs in the original report [9] coincided with B cell DVSs identified in this analysis, while only 5% (55/1033) of T cell DMCs were overlapped with T cell DVSs. It suggests that higher variation in B cell methylation might be a major reason for the low number of DMRs identified in B cells.

### PPI network analysis revealed a hypermethylated TCR signaling cluster and a BCR signaling cluster with highly variable methylation

To understand the functional implications of these DMRs and DVSs, we performed further analyses focusing on the genes with promoter-region DMRs or DVSs, respectively. Through gene ontology analysis by DAVID [28], we found significant enrichment in genes involved in innate immunity for the hypomethylated genes in T cells, typically related to the type I interferon pathway. In contrast, hypermethylated genes were found predominantly involved in adaptive immunity, especially in T cell receptor (TCR) signaling (Additional file 2: **Supplementary Fig. 2a**). Interestingly, we found strong enrichment of B cell receptor (BCR) signaling for the hypervariable genes in B cells, as indicated by KEGG pathway analysis (Additional file 2: **Supplementary Fig. 2b**).

Furthermore, using network analysis based on protein-protein interaction data (PPI) implemented in the XGR software [24], we obtained a maximum-scoring subnetwork of DMRs (**Fig. 2**) from the STRING database of functional PPIs [25]. Within this subnetwork, two gene modules were most prominent from the rest of the network, one dominated by hypomethylated (green) and the other by hypermethylated (red) genes. These modules further supported the finding above that the hypo- and hypermethylated genes clustered largely separately in the network and were enriched in innate- (Bonferroni-adjusted P-value = 6e-26) and adaptive immunity (Bonferroni-adjusted P-value = 4e-06), respectively.

**Fig. 2.**
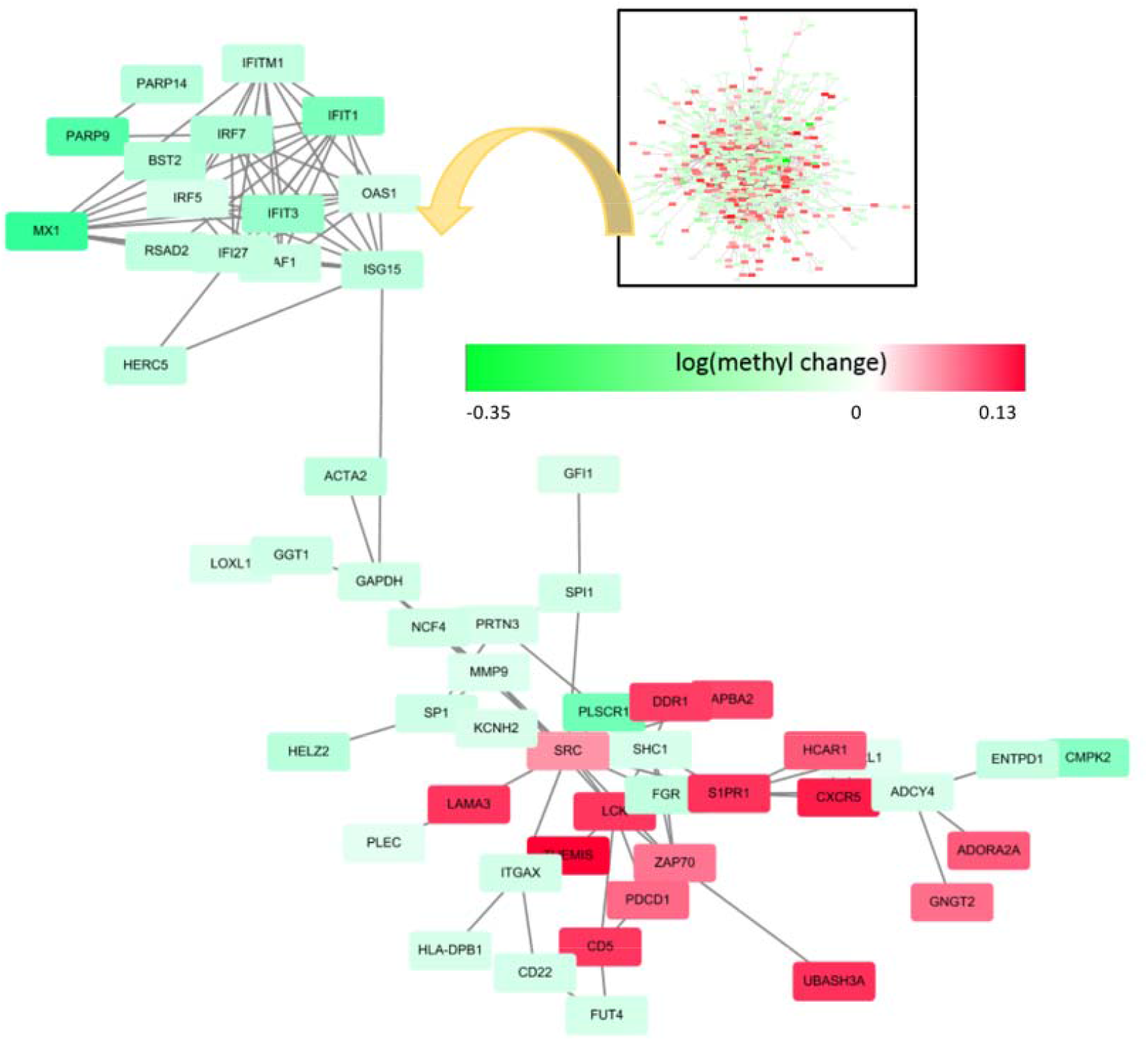
The two most outstanding modules in a PPI network analysis of DMRs reveal a hypomethylated innate immunity gene cluster and a hypermethylated adaptive immunity gene cluster. The optimal PPI subnetwork containing most highly significant DMRs in T cells, extracted from the overall PPI network using all T cell DMRs (**inset**). A hypermethylated gene cluster (red color) were enriched in TCR co-signaling, while the hypomethylated cluster (green) were mainly enriched in Type I interferon signaling.

Functional annotations by pathway analysis were also performed for all the genes in this subnetwork (Additional file 2: **Supplementary Fig. 3a**). They were found particularly enriched in PD-1 signaling, a co-inhibitory pathway to T cell activation (FDR < 0.01). Key genes in this pathway, as curated in the Reactome Pathway Database, including *PDCD1* (i.e. *PD-1*), *LCK, CD247, CD3D, CD3E*, and *CD3G*, were all found hypermethylated, as well as other inhibitory immune checkpoint molecules, such as *CTLA4* and *BTLA*. Furthermore, many T cell receptor (TCR) co-stimulatory molecules, including *CD28, CD2*, and *SLAM* were also found hypermethylated. All the hypermethylated molecules involved in TCR co-signaling (stimulatory or inhibitory [29]) were shown in **Fig. 3**. Using the same procedure of PPI subnetwork analysis, we observed a hypervariable methylation cluster in B cells, which was significantly enriched in the BCR signaling pathway (Additional file 2: **Supplementary Fig. 3b-c**).

**Fig. 3.**
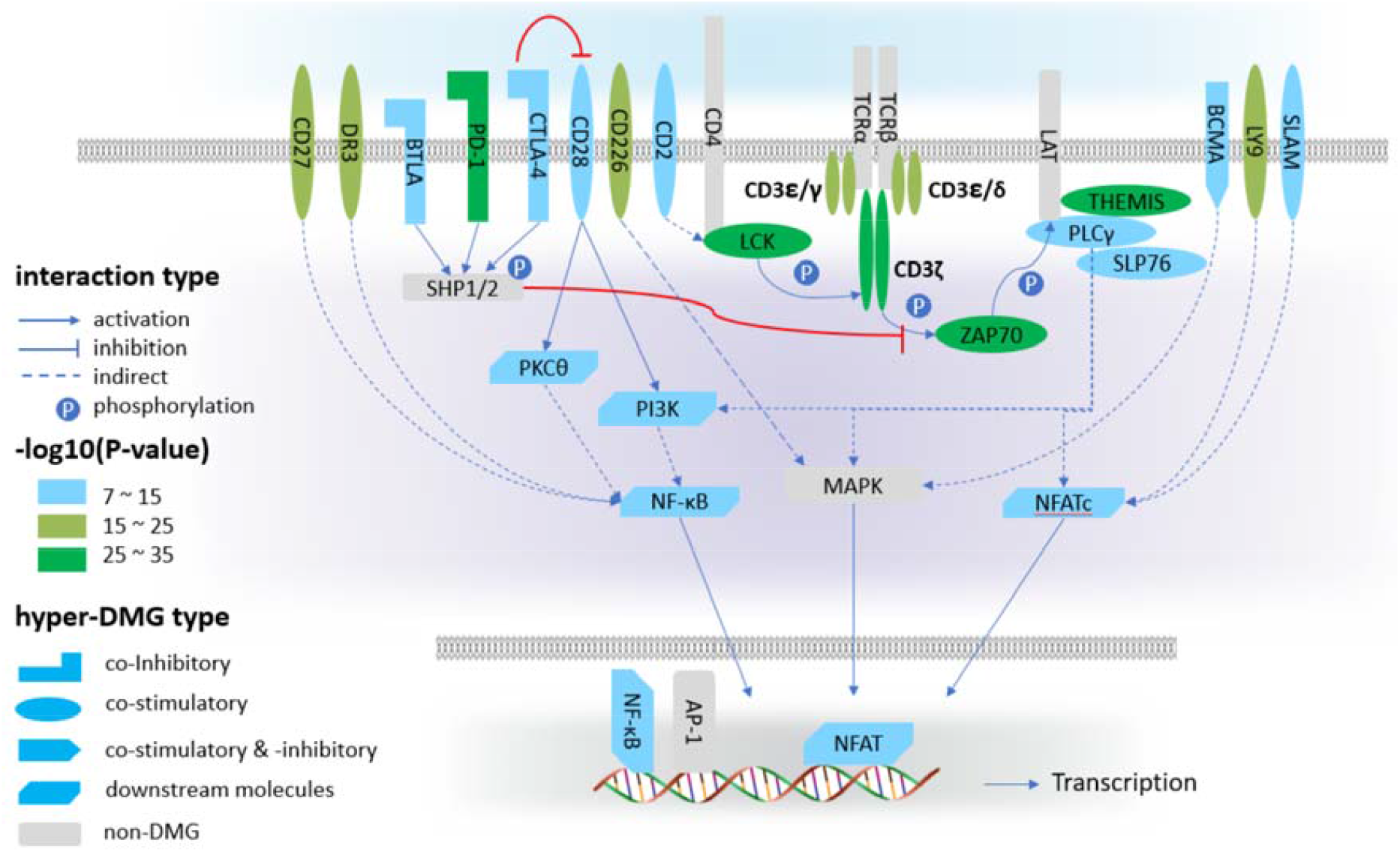
Hyper-methylation of T cell receptor (TCR) co-signaling molecules. All colored shapes indicate hyper-methylated genes and grey shapes stand for genes without significant methylation change in SLE patients.

### DMRs and DVSs and their corresponding chromatin states

Chromatin states are defined by histone modifications, including histone methylation (H3K4me1, H3K4me3, H3K9me3, H3K27me3, and H3K36me3) and acetylation (H3K27ac) [30]. The genomic sequences are categorized as TSS, transcription, enhancer, bivalent, and repressed regions. Using the same enrichment method for the suggestive GWAS loci (Additional file 2: **Supplementary Results**), we determined the over- and under-representation of DMRs and DVSs in different chromatin states (Additional file 1: **Tables S8**). Expectedly, we found strong enrichment of DMRs in TSS regions and depletion in repressed PolyComb and quiescent regions (**Fig. 4a**), with similar patterns between hypermethylated and hypomethylated DMRs. Interestingly, for hypervariable regions in B cell DVSs, we observed strongest enrichment in enhancers and depletion in active TSS regions (**Fig. 4b**).

**Fig. 4.**
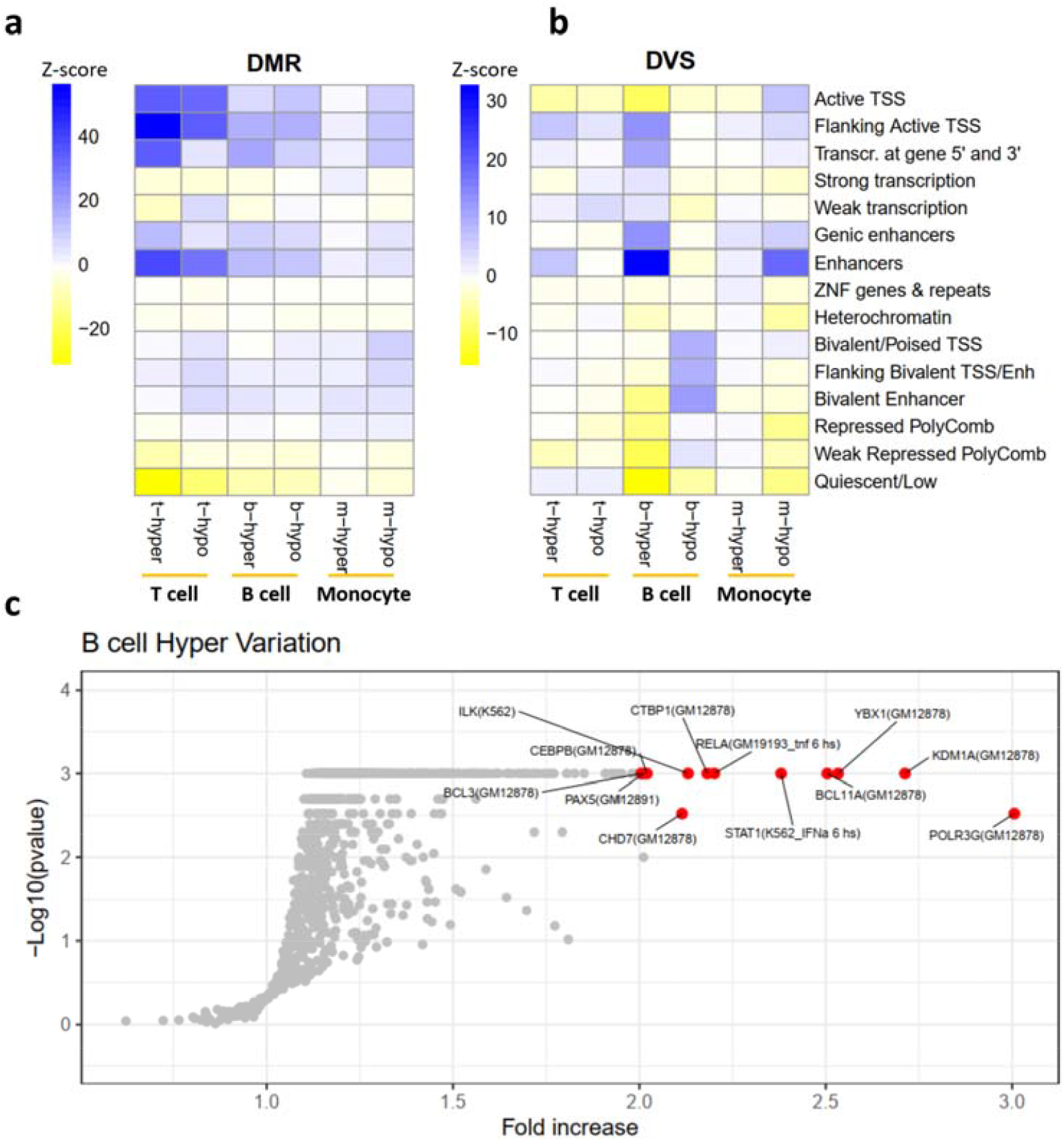
DVSs were enriched in enhancer chromatin states and binding sites of enhancer-related TFs, as determined by permutation test. (**a-b**) For enrichment of chromatin states, TSS regions had the strongest enrichment in DMRs in both B cells and T cells, and repressed PolyComb regions were depleted (**a**). Enhancer regions had the strongest enrichment in B cell DVSs. Active TSS regions were particularly depleted in DVSs across cell types (**b**). (**c**) TF enrichment in hypervariable DVSs in B cells. Only several top-ranked TFs were labeled for visualization. Several enhancer-binding TFs were overrepresented in hypervariable DVSs in B cells, including KDM1A, CEBPB, CHD7, and POLR3G.

### Transcription factors enriched in SLE DMRs and DVSs

TFs are key elements for gene expression regulation. Here we investigated the TFs bound to the proximal promoter regions of the genes with SLE DMRs or DVSs (Additional file 2: **Supplementary Fig. 4-5**, Additional file 1: **Tables S9-12**). Interestingly, for DVSs in B cells, we found enrichment of TFs with the ability to bind enhancers (EP300 [31] and CEBPB [32]), which may have connection with the overrepresentation of enhancer chromatin for hypervariable DVSs in B cells (**Fig. 4b**). We found several other top-ranked enriched TFs that were also reported to be able to bind enhancers, including KDM1A [33], CHD7 [34], PAX5 [35], and CTBP1 [36] for the hypervariable DVSs in B cells (**Fig. 4c**). Among these TFs, PAX5 is a B-cell specific activator that is specifically expressed at the early stage of B-cell differentiation. CEBPB is important in immune and inflammatory responses, such as differentiation of B cells and macrophages [37]. Their interactions in PPI networks were shown in Additional file 2: **Supplementary Fig. 6**. Of note, many of the highly-ranked TFs were from experiments performed by using lymphoblast like cell lines, such as GM12878 and K562, which may reflect the cell-specificity of the DMRs.

### DMR genes distinguished patients from controls and predicted disease activities

An interesting question is whether SLE patients can be distinguished from healthy individuals based on the methylation states of CpG sites or genes. To this end, we applied a machine learning algorithm, LASSO (Least Absolute Shrinkage and Selection Operator) [38], using 10-fold cross-validation. We identified 44 CpGs from 22,550 of those within the T cell DMRs. We constructed a generalized linear model based on the T cell methylation densities of these 44 CpGs (**Fig. 5a)**, and applied this model to all datasets for SLE across various cell types, including T cells, B cells, monocytes, and PBMCs. As shown in **Fig. 5b-c**, this model achieved high accuracy across different cell types for SLE, with the area under the receiver operating characteristic (ROC) curve (AUC) reaching about 0.990 (**Fig. 5c**). Similarly, training the model based on CpG sites in the B cell DMRs with the same procedure identified 36 CpGs, which achieved similar prediction accuracy for SLE (AUC 0.980) (Additional file 2: **Supplementary Fig. 7a-b**). Including DVSs in the models did not further improve their performance.

**Fig. 5.**
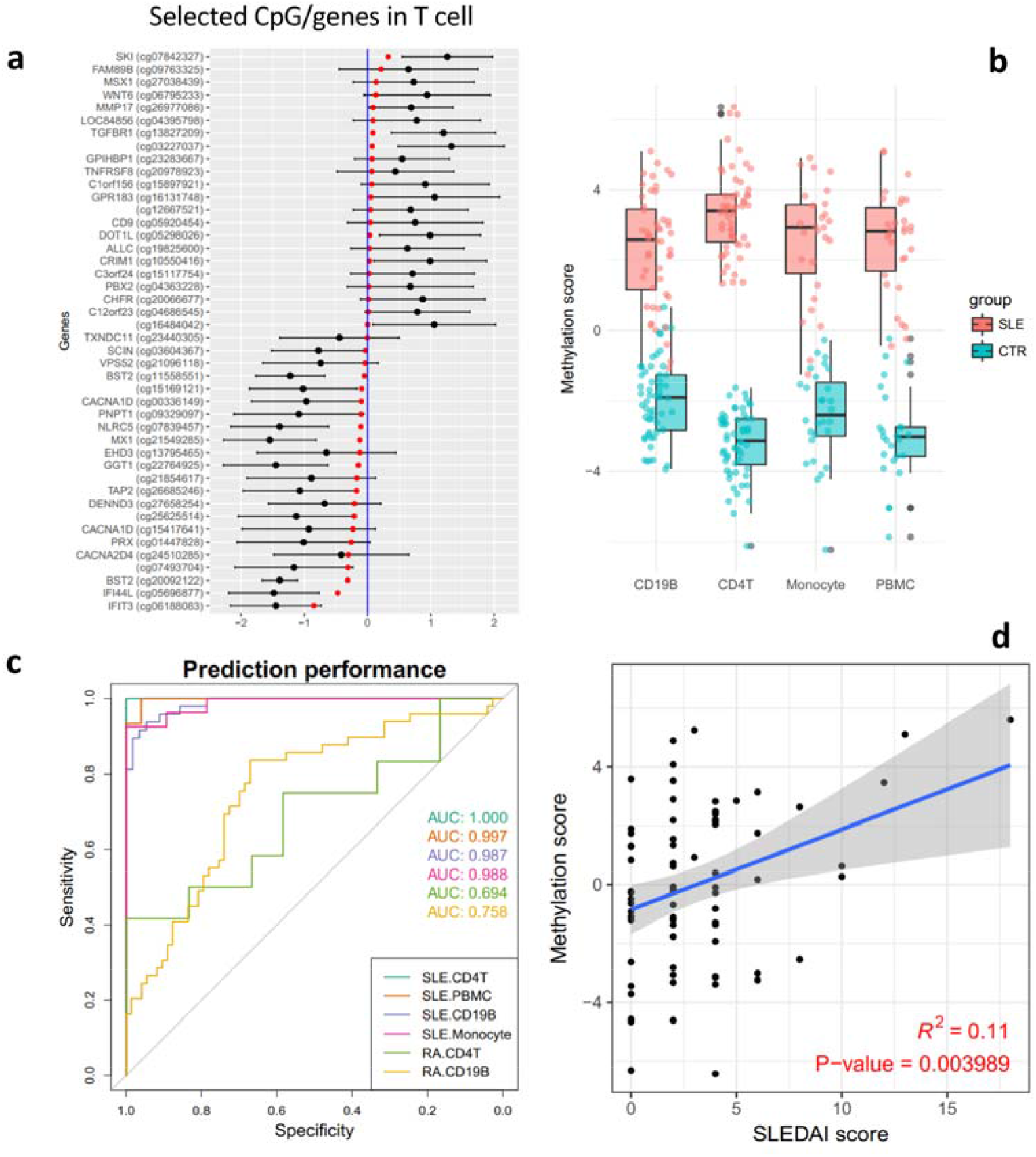
Machine learning identified DMR genes accurately distinguishing patients from controls and correlated with SLE disease activity. (**a**) A list of DMR genes in T cells were selected by LASSO with different weights. Red dots indicate model coefficients and black dots with error bars show the normalized methylation difference between patients and controls for CpGs. (**b**) Performance of the trained model by T cell data (shown in **a**) in different datasets, as defined by separation of SLE patients from controls. (**c**) Model performance of (**a**) measured by AUC, with very good values for each cell type in SLE (0.97 on average) but much lower in RA (0.72 on average). (**d**) Significant correlation between SLEDAI scores and M-scores for SLE patients, using naïve CD4+ T cell samples (R = 0.33).

Collectively, the 75 selected CpGs from T cell (44 CpGs) and B cell DMRs (36 CpGs) were mapped to a total of 60 genes, nine of which were reported to be differentially methylated in naïve CD4+ T cells of SLE patients in another study [12]. Among them, methylation of *IFI44L* in PBMC has been reported as a biomarker for SLE with high sensitivity and specificity [39], although a different CpG site in the gene was identified in this study. Interestingly, clear separation of SLE patients from controls by the reported CpGs in *IFI44L* only worked for PBMC samples, but not as well for samples from T cells, B cells, or monocytes (Additional file 2: **Supplementary Fig. 8**).

Functional enrichment analysis of these 60 genes showed over-representation of type I interferon signaling (Additional file 2: **Supplementary Fig. 9a**). In addition to the overall DMR enrichment in SLE GWAS risk loci (Additional file 2: **Supplementary Results**), we found that these genes provided further information on disease mechanisms when analyzed together with the susceptibility genes. Importantly, pathway analysis identified significant role of phagosome pathway (Bonferroni-adjusted P-value=0.028) in the disease after including DMR genes *HLA-B* and *TAP2* (Additional file 2: **Supplementary Fig. 9b**) in the analysis.

Defects in this pathway were previously reported to be associated with SLE [40]. PPI network analysis showed that 24 of the 60 DMR genes had interactions with susceptibility genes, far more than expected by chance (odds ratio, OR =2.2, Chi-squared test P-value <1e-16), using a method published recently [41]. Further PPI clustering analysis identified modules including T-cell co-stimulation and calcium signaling (Additional file 2: **Supplementary Fig. 10, Supplementary Methods**).

Finally, we investigated whether there are correlations between methylation scores (M-score), defined as the weighted sum of methylation of the selected CpGs in T cells or B cells, respectively, and clinical manifestations such as age, ethnicity, and SLEDAI disease activity score. Using a recent published dataset on DNA methylation in naïve CD4+ T cells from SLE patients [10]. we found no significant correlation between M-score in T cells and ethnicity or age (Additional file 2: **Supplementary Fig. 7c-d**). However, we observed a significant correlation of M-score with SLEDAI scores (**Fig. 5d**, Pearson correlation R-squared = 0.11, P-value = 0.004). Similar results were observed for M-scores in B cells (Additional file 2: **Supplementary Fig. 7e**).

## Discussion

In this study, we detected differentially methylated regions and methylated sites with differential variability in SLE patients and analyzed their implications in SLE pathogenesis. We found most DVSs in B cells while the majority of DMRs were in T cells, enriched in the BCR and TCR signaling pathway, respectively. DVSs were specifically enriched in enhancers and depleted in active promoters, and they were also significantly enriched in the binding regions of several enhancer-binding TFs, including EP300, CEBPB, KDM1A, CHD7, and PAX5. Although we have developed an efficient model predicting SLE with high sensitivity and specificity based on a group of informative genes, much work is needed to characterize them for clinical use in the future.

As an important form of epigenetic modifications, DNA methylation is closely related to and may fluctuate with changes of cellular environment. Signals from single CpG based analysis were prone to false positives due to hidden confounding factors and biological variations. In contrast, region-based detection of differential methylation has higher sensitivity, without sacrificing specificity [17]. The smoothing technique in region-based analysis made it more robust to outliers, as it considers strong correlations among adjacent CpG sites [26]. In addition, CpG-based analyses suffer more on multiple-testing burden and also may identify different CpG sites that are highly correlated, which are more likely to be reported as a single DMR by region-based analysis. For example, in the original report [9], cg12044210 and cg21917349 for gene *APBA2* were both reported as differentially methylated CpGs, but they are only 2bp away from each other and are highly correlated (Pearson correlation 0.92).

Probably due to the higher sensitivity of DMR analysis, we identified a number of genes as hypermethylated in T cells, which may have been alluded to in the original study by Absher et al. (Figure 2B [9]). They were not explored in any detail, as the Absher study adopted an extremely stringent P-value threshold and as a result, most of the CpGs surpassing their P-value threshold were those involved in the Type I interferon signaling pathway. Network analysis based on protein-protein interaction information revealed a hypermethylated gene module predominantly involved in T-cell receptor co-signaling (**Fig. 2**). These findings expanded our understanding of this prototype autoimmune disease compared to the widespread focus on the hallmark type I interferon signature by most epigenomic or transcriptomic studies for SLE. We have no information on whether the patients were under treatment of immunosuppressive drugs at the time of sample collection and cannot rule out the possibility that the hypermethylated changes in TCR signaling molecules was an effect of immunosuppressive treatment. On the other hand, it was recently reported that drugs mostly decreased DNA methylation in patients of SLE [16].

Half of the DMRs identified in our analysis were replicated in an independent dataset generated from PBMC using the same analysis procedures [11]. We also compared our DMRs to 7,625 differentially methylated CpGs identified from a recent study using whole blood from SLE patients [16]. In T cells, we found that 41% (337) of the DMRs with at least one significant CpG in the original report by Absher et al. [9] also had differentially methylated CpGs reported by Imgenberg-Kreuz et al. In contrast, 46% (1,325) of the novel DMRs we identified in this study had at least one CpG that is significant in the study by Imgenberg-Kreuz et al. It suggested that the novel DMRs detected from this analysis should have a comparable false positive rate compared to the DMRs containing significant differentially methylated CpGs in the original study.

We observed big difference in the numbers of DMRs between T cells and B cells and thought that alternative analysis other than on the differences in average methylation levels may reveal more information on methylation aberrations. Interestingly, we found that there were more methylation variable sites in B cells of SLE patients than in T cells (Additional file 2: **Supplementary Fig. 1a**), which may explain the fewer number of DMR sites found in B cells. Highly variable methylation in SLE B cells is an interesting phenomenon and was not reported before to the best of our knowledge. It might reflect a high-level heterogeneity in B cells among patients, something worth further exploration in future studies, such as potential correlations with clinical subphenotypes. Genes involved in BCR signaling pathway were found to be particularly variable in their methylation level among patients. Our study highlights the need to further analyze public available methylation data to gain additional insights in disease mechanisms.

## Conclusions

Through novel analysis of methylation data, we identified abnormal hypermethylation in genes involved in TCR co-signaling pathways, and higher methylation viability in BCR signaling pathways. We showed that DVSs were specifically enriched in enhancers, which was supported by a group of enhancer-binding TFs enriched in genes with DVSs, suggesting an important role of an enhancer-mediated regulatory network in SLE. Finally, we used LASSO, a machine learning approach, to select a list of CpG sites that accurately predicted SLE disease status, and also correlated with disease activities. These findings may help us better understand the mechanisms of this prototype autoimmune disease, and to develop novel biomarkers and treatment targets.

## Supporting information

Supplementary Results and Figures

## Additional Files

Additional file 1: **Table S1**. DMRs in T cells. **Table S2**. DMRs in B cells. **Table S3**. DMRs in monocytes. **Table S4**. DMRs in PBMCs. **Table S5**. DVSs in T cells. **Table S6**. DVSs in B cells. **Table S7**. DVSs in monocytes. **Table S8**. Chromatin state enrichment in DMRs and DVSs. **Table S9**. TF enrichment in T cell DMRs. **Table S10**. TF enrichment in B cell DMRs. **Table S11**. TF enrichment in T cell DVSs. **Table S12**. TF enrichment in B cell DVSs. **Table S13**. DMG bound by STATs. **Table S14**. GWAS loci with DMRs or DVSs

Additional file 2: **Supplementary Results**. Additional results. **Supplementary Methods**. Additional methods. **Supplementary Fig. 1. Comparison between DVSs and DMRs or DMCs. Supplementary Fig. 2. Functional enrichment for T cell DMRs and B cell DVSs. Supplementary Fig. 3. TCR and BCR signaling clusters in the PPI subnetwork. Supplementary Fig. 4. TF enrichment in DMRs. Supplementary Fig. 5. TF enrichment in DVSs. Supplementary Fig. 6. Interactions of enhancer-binding TFs enriched in hypervariable DVSs in B cells. Supplementary Fig. 7. Machine learning and correlation of prediction scores with clinical variables. Supplementary Fig. 8. DNA methylation of two CpG sites in IFI44L in SLE and controls across different cell types. Supplementary Fig. 9. Functional enrichment analysis for DMR genes selected by LASSO. Supplementary Fig. 10. PPI network modules defined by GWAS risk genes and DMR genes selected by LASSO. Supplementary Fig. 11. Examples of DMRs and DVSs identified in known SLE susceptibility loci. Supplementary Fig. 12. Additional representative known SLE associated genes with DMRs. Supplementary Fig. 13. Examples of DMRs and DVSs identified in known SLE susceptibility loci. Supplementary Fig. 14. Enrichment of the suggestive GWAS loci (Pvalue > 5e-8 and < 1e-3) in DMRs. Supplementary Fig. 15.Enrichment of suggestive GWAS signals (P-value > 5e-8 and < 1e-3) in DVSs. Supplementary Fig. 16. STAT1 and STAT2 upon stimulation were differentially enriched in hypomethylated DMRs in B cells. Supplementary Fig. 17. Correlation between DNA methylation and gene expression in SLE. Supplementary Fig. 18 Three examples of DMR, DVS, and DMC in SLE T cells using UCSC Genome Browser**.

## Abbreviations

AUC: Area under the receiver operating characteristic curve
BCR: B cell receptor
DMC: Differentially methylated CpG
DMR: Differentially methylated region
DVS: Differentially variable site
FDR: False discovery rate
GEO: Gene expression omnibus
GWAS: Genome-wide association study
LASSO: Least absolute shrinkage and selection operator
PBMC: Peripheral blood mononuclear cells
PPI: Protein-protein interaction
ROC: Receiver operating characteristic
RA: Rheumatoid arthritis
SLE: Systemic lupus erythematosus
SLEDAI: Systemic lupus erythematosus disease activity index
TF: Transcription factor
TFBS: Transcription factor binding sites
TSS: Transcription start site
TCR: T cell receptor

## Declarations

### Ethics approval and consent to participate

Not applicable

### Consent for publication

Not applicable

### Availability of data and material

All data generated or analysed during this study are included in this published article and its supplementary information files.

## Acknowledgements

The authors thank Hong Kong PhD fellowship scheme (HKPF), HKU Postgraduate Scholarships, and the Edward & Yolanda Wong Fund for supporting postgraduate students who participated in this work.

## Authors’ contributions

WY, YLL, and MG proposed and carried out the research and wrote the manuscript. TYW and YFW contributed to the TF analysis. JJS contributed to statistical analysis and edited the manuscript. All authors read and approved the manuscript.

