## Supplementary Results and Figures for "Region- and variance-based DNA methylation analyses reveal novel disease genes and pathways for systemic lupus erythematosus"

**In-depth analysis of differential DNA methylation reveals novel disease genes and pathways in SLE**

### **Supplementary Results**

#### **Comparing DMRs in this analysis with significant DMCs from the original report**

We found 96% (993 out of 1,033) of the significant CpGs reported in the original publication on this dataset [1] were also found in 807 of the 3,720 T cell DMRs identified in this study. The numbers of DMRs with or without originally reported CpGs is shown in **Fig. 1c**. More than three fourths of the DMRs were novel from this study, largely detected from T cells, which was expected given the advantages of region-based statistical models over CpG-based methods [2]. Single-CpG analyses often had to adopt more stringent P value thresholds because they are more prone to false positives, due to hidden confounding factors and variations on a single CpG site. In addition, CpG-based analyses suffer more on multiple-testing burden and also may identify different CpG sites that are highly correlated, which are more likely to be reported as a single DMR by region-based analysis. For example, in the original report, cg12044210 and cg21917349 for gene *APBA2* were both reported as differentially methylated CpGs, but they are only 2bp away from each other and are highly correlated (Pearson correlation coefficient of 0.92). In fact, a total of 291 (>25%) originally reported differential CpGs were within 500 bp of each other and might be correlated with each other. In addition, we found that 26% (44/166) of the B cell DMCs in the original report were among the B cell DVSs identified in this analysis, while only 5% (55/1033) of T cell DMCs were overlapped with T cell DVSs. It suggests that higher variation in B cell methylation might be a major reason for the low number of DMRs identified in B cells.

#### **Validation of DMRs in PBMCs**

To validate DMRs identified in T cells, B cells, and monocytes, we applied the same DMR calling method on an independent dataset generated on PBMC from SLE patients and controls of Chinese ancestry (Additional file 1: **Table S4**). Applying the same significance threshold of 1.0e-7, we identified a list of 7,230 (4,121 or 57% hypomethylated and 3,109 or 43% hypermethylated) DMRs [3]. Although both the cell types and the ethnic origins were different between the two datasets, high level overlapping for the hypomethylated DMRs was observed between PBMCs in the Chinese study and T cells, B cells and monocytes in the European cohort, namely 1,242 (53%), 103 (56%), and 47 (70%) for the three cell types, respectively. The overlaps for hypermethylated DMRs were much lower, at 569 (41%), 35 (19%), and 10 (24%) for T cells, B cells, and monocytes, respectively. DMRs with discordant directions in the two datasets were found to be high in B cells, 64 (17%) but not for T cells and monocytes, 117 (3%) and 7 (6%), respectively, probably reflecting smaller fractions of B cells in PBMC and higher variance for B cell methylation changes.

#### **Differential enrichment of transcription factors in SLE DMRs and DVSs**

We investigated the TFs bound to the proximal promoter regions of the genes with SLE DMRs or DVSs, respectively, in various cellular conditions based on ENCODE data. Overall, we found that STATs (STAT1, STAT2, and STAT3) and IRFs (IRF1 and IRF9) were particularly enriched in SLE hypomethylated genes. STAT1 is known to dimerize with STAT2 to form ISGF3 complex. Together with IRF9, they stimulate the expression of target genes with interferon-stimulated response elements (ISRE), such as OAS2, MX1 and IFIT1 [4] (Additional file 2: Supplementary Fig. 4**,** Additional file 1: **Tables S9-10**). Additionally, we observed enrichment of EZH2 in hypermethylated DMRs in T cells, a result consistent with previous report (Additional file 2: Supplementary Fig. 4**b**) [5]. Moreover, we observed enrichment of EZH2 in DVSs in both T cells and B cells (Additional file 2: Supplementary Fig. 5**,** Additional file 1: **Tables S11-12**).

#### **Additional functional annotation for the selected DMR genes by LASSO**

Some of the DMR genes selected by LASSO play important roles in immune-related pathways. The first one is TGF-β signalling. *TGFBR1* encodes a type-I TGF-β receptor that binds TGF-β and transduces extracellular signals to regulate cellular processes, such as cell differentiation and apoptosis. SKI cooperates with FAM89B to negatively regulate TGF-β signalling by preventing the translocation of SMAD2 from nucleus to cytoplasm in response to TGF-β, which is critical for activating the regulatory T cells [6]. Other pathways include NF-κB and JAK-STAT signalling. *TNFRSF8* (*CD30*) encodes a tumour necrosis factor (TNF) receptor which is only expressed on activated T- and B cell surface. It can lead to the activation of NF-κB and help prevent autoimmunity [7, 8]. DOT1L was also reported to interact with STAT1 and regulate *IRF1* expression, thus involved in the JAK-STAT signalling pathway [9].

A number of DMR genes selected by LASSO are reported to be associated with SLE pathogenesis by either genetics or epigenetics studies. *TNXB*, localized in the MHC class III region, encodes for a member of the extracellular matrix glycoproteins called tenascins. This protein has anti-adhesive effect, opposite to the adhesive effect of integrin proteins. Both *TNXB* and integrin subunits, such as *IGTAL* and *ITGAM*, were reported to be involved in SLE pathogenesis, and ITGAM was even one of the strongest genetic risk factors for SLE [10-13]. Moreover, B cell receptor (BCR) signaling and autoantibody generation were closely related to SLE. PIK3R5 participates in signaling processes including cell proliferation and survival. It was hypomethylated in SLE B cells in our analysis and reported to be remarkably up-regulated together with USP18 in the BCR pathway upon stimulation of IFNα and anti-IgM [14]. The hypomethylation of *PARP9* and *PARP14* was associated with auto-antibody generation in lupus patients [15]. *DUOX2* (Dual Oxidase 2) is a subunit of complex generating hydrogen peroxide (H_2_O_2_), which is required by thyroid peroxidase (TPO) activity, to which autoantibodies were found in SLE patients and associated with disease activity [16, 17].

#### **Evaluate disease specificity of the LASSO model trained on SLE DMRs by applying it to rheumatoid arthritis**

We asked whether this model, derived from SLE, also applied to rheumatoid arthritis (RA), an autoimmune disease that shared a number of susceptibility genes with SLE. Although RA samples showed slightly higher prediction score on average than healthy controls, we do not find good separation between RA samples and controls (Additional file 2: Supplementary Fig. 7**f**), either for CD4 T cells or for CD19 B cells [18, 19], and the AUC for RA were much lower than that of SLE (Additional file 2: **Fig. 5c,** Supplementary Fig. 7**b**).

#### **PPI clustering analysis of GWAS susceptibility genes with the selected DMR genes by LASSO**

To functionally annotate the 60 genes whose methylation accurately distinguished SLE patients and controls, we performed gene ontology (GO) analysis on them. The analysis showed over-representation of type I interferon signaling and defense response to virus (Additional file 2: Supplementary Fig. 9**a**). Since we demonstrated that DMRs were highly enriched in SLE GWAS risk loci, here we asked whether the 60 selected genes would provide further information in explaining disease mechanisms when analyzed together with the susceptibility genes. We analyzed the pathways and PPI information on these differentially methylated genes together with the about 90 reported SLE risk genes. Interestingly, we found that some pathways were improved or became significant only after integrating GWAS genes with the DMR genes (Additional file 2: Supplementary Fig. 9**b**), such as herpesvirus infection pathway and cytokine-cytokine interaction pathway. Importantly, the phagosome pathway became significant (Bonferroni-adjusted P-value=0.028, nominal P-value=0.004, compared with GWAS-only Bonferroni-adjusted P-value=0.068, nominal P-value=0.018) after including *HLA-B* and *TAP2* from DMRs. Defects in this pathway were reported to be associated with SLE previously [20].

Apart from GO and pathway enrichment, we tried to analyze protein interaction between GWAS risk genes and the 60 DMR genes. Using PPI network from STRING database [21], we observed that 24 of the 60 DMR genes (40%) had interactions with GWAS risk genes, far more than expected by random chance (odds ratio, OR =2.2, Chi-squared test P-value <1e-16), by adopting a method published recently [22]. In this analysis, we confirmed the significant interaction of targets of SLE drugs and risk genes (OR =2.9, Chi-squared P-value <1e-16) and found significant interaction between these DMR genes and SLE drug targets as well (OR=1.4, Chi-squared P-value <1e-16).

Furthermore, within this PPI network, we investigated potential modules formed by GWAS risk genes and the 60 DMR genes. We identified eight PPI modules in total, with the largest one composed of only GWAS risk genes involved in regulation of innate immune response (Additional file 2: Supplementary Fig. 10a). Of note, DMR gene *PIK3R5*, a kinase important in cell proliferation and survival, formed a cluster with six other GWAS risk genes related to leukocyte proliferation. In contrast, SLE susceptibility gene *HIP1*, a proapoptotic protein, clustered with five DMR genes involved in response to type I interferons. Among the other clusters were those related to T-cell co-stimulation, macroautophagy, and protein SUMOylation (Additional file 2: Supplementary Fig. 10a).

Interestingly, we identified a dense four-node module (each gene interacted with all other genes within the module) containing two DMR genes, *CACNA1D* and *CACNA2D4*, and two GWAS risk genes, *RASGRP1* and *RASGRP3*. These four genes were important in linking calcium signaling with MAPK pathway (Additional file 2: Supplementary Fig. 10b), both of which were known to be involved in SLE [23, 24]. Moreover, by searching public gene expression dataset (GSE21546) [25], we found that 13 of the 60 DMR genes were upregulated (FDR<=0.05) after knocking out *ELK4* in double-positive thymocytes (enrichment FDR= 3e-12) [25]. suggesting a key role of *ELK4* in the pathogenesis of the disease. *ELK4*, which is phosphorylated by *MAPK1* and *MAPK8* (Mitogen-activated protein kinase) [26, 27], is a member of ETS family transcription factor and is required for thymocyte positive selection.

#### DMRs and DVSs were enriched in loci associated with SLE

DNA methylation may act as a mediator for genetic risk factors to affect disease phenotypes, or the genes involved in disease pathogenesis may have both genetic and epigenetic aberrations. For the susceptibility loci reaching genome-wide significance [28], we found significant enrichment of DMRs (hypergeometric P-value = 6e-10 for T cells and P-value = 9e-5 for B cells, Additional file 1: Table S8) and DVSs (P-value = 5e-6 for B cells, and insignificant for T cells, Additional file 1: Table S8). Notable DMR examples include *IRF7*, *ETS1*, *CD247,* and *BLK*, the latter three of which were hypermethylated and *IRF7* was hypomethylated (Additional file 2: Supplementary Fig. 11), and more examples were shown in Additional file 2: Supplementary Fig. 12.

*IRF7* had the largest differences between cases and controls genome-wide, and hypomethylation occurred across the entire gene as well as its up- and down-stream regions (Additional file 2: Supplementary Fig. 11a) in all three cell types, with monocytes having the most dramatic differences. Expression level of *IRF7* was found to be upregulated significantly in SLE samples, namely by 3-fold (FDR= 0.003) in T cells, and by 2-fold (FDR= 0.003) in B cells (Wang et al., manuscript under review). *IRF7* is a transcription factor involved in the expression of type I interferon genes (IFN) and interferon-stimulated genes (ISG), both of which are hallmarks of innate immunity.

Previously, *ETS1* risk alleles were found to be associated with lower expression of the gene and were correlated with increased levels of IL-17 in the peripheral blood [29, 30]. Hypermethylation of this gene (Additional file 2: Supplementary Fig. 11b) might be one of the mechanisms hindering its role as a negative regulator of B cell and Th17 cell differentiation in SLE. *CD247* encodes a subunit of the CD3 protein complex, which forms the TCR-CD3 complex. It functions almost exclusively in T cells, which is reflected by the chromatin states in this region in various cell types [31] (Additional file 2: Supplementary Fig. 11c). Hyper-methylation in *CD247* was also reported previously in SLE [32]. *BLK* is a B cell tyrosine kinase, which is important for B lymphocyte development, was found hypermethylated in both T and B cells (Additional file 2: Supplementary Fig. 11d). Like *ETS1*, reduced expression of *BLK* was known to be associated with SLE [33]. Additionally, in Additional file 2: Supplementary Fig. 13a-b, we have showed that *ETS1* and *BLK* were generally hypermethylated but with high variation in patient B cells but not in control samples, which was detected by DVS analysis. In T cells, *ETS1* and *BLK* had comparable variance between patients and controls, but highly significant DMR P-values than in B cells (Additional file 2: Supplementary Fig. 13c-d).

We also observed significant enrichment of DMRs in the suggestive SLE loci with genetic association P-value < 1e-3 and > 5e-8. Suggestive variants, 27,644 in total, were identified from our published three-way meta-analysis of the genome-wide association studies from UK, Hong Kong and Anhui (China) [34]. We extended 50kb upstream and downstream around the associated variants and merged overlapping regions to obtain a total of 1,754 suggestive risk loci. The observed overlap between DMRs and suggestive GWAS loci was significantly larger than the size- and microarray background-matched reshuffled DMR regions (z-scores were 9.0, 7.4, and 3.5 for T cells, B cells, and monocytes, respectively, Additional file 2: Supplementary Fig. 14). Interestingly, we observed a significant DVS enrichment in suggestive loci for B cells, but not for T cells (z-scores were 0.3, 4.2, and 2.9 for T cells, B cells, and monocytes, respectively, Additional file 2: Supplementary Fig. 15).

A recent study by Imgenberg-Kreuz et al. found link between DNA methylation and susceptibility variants, reporting seven methylation quantitative trait loci (meQTL) that are also SLE risk variants, such as *IRF7* and *UBE2L3*. Thus, it is possible that some genetic risk loci may exert their effect on disease risk through altering DNA methylation, although it is also possible that they are involved in the disease through mechanisms different from methylation. Compared to Illumina 450k BeadChip platform, whole genome bisulfite sequencing (WGBS) might be better suited to more systematically search for the link between genetic and epigenetic changes.

#### Differential enrichment of transcription factors upon interferon stimulation in SLE DMRs

In ENCODE, a few TF ChIP-Seq experiments were conducted under stimulation of interferons for a different period, which provided an opportunity to study the role of interferons on TF binding to the genes with SLE DMRs. In general, we found a number of TFs with greater enrichment upon longer period of interferon stimulation, largely through binding to a larger number of differentially methylated genes (DMG), defined as those genes with DMRs in the promoter-proximal region of 2 Kb up- or downstream of their TSS. Take hypomethylated DMGs in B cells as an example. For the K562 cell line, a human immortalized myelogenous leukemia cell line, STAT2 had 17-fold enrichment upon IFN alpha (IFNα) stimulation for 6 hours (STAT2_IFNα_6h) compared to just 4.8-fold enrichment with 30-minute stimulation (STAT2_IFNα_30min) (Additional file 2: Supplementary Fig. 16a, Additional file 1: Table S13). In contrast, STAT1 only had a mild increase of enrichment – from 14-fold for 30-minute stimulation (STAT1_IFNα_30min) to 18-fold for 6-hour treatment of IFNα (STAT1_IFNα_6h) (Additional file 2: Supplementary Fig. 16b). Moreover, with the same duration of treatment, IFNα (STAT1_IFNα_6h, 18-fold) seems to have larger effect than IFNγ (STAT1_IFNγ_6h, 3.3-fold) on *STAT1*, which is consistent with the more important role of STAT1 in IFNα signaling (Additional file 2: Supplementary Fig. 16c). However, large fold changes of STAT1 or STAT2 upon stimulation were not observed in DVSs, which may suggest a lack of an interferon component in DVSs, an underlying difference from DMRs.

After looking into the DMGs bound by STAT1 or STAT2 with interferon treatments, however, we did not find any significant increase of the number of TF binding sites per gene. Yet 10 of the 22 DMGs only appearing in STAT1_IFNα_6h dataset had either physical interaction (based on PPI data) or colocalization to the same subcellular compartment with those DMGs in STAT1_IFNγ_6h (Additional file 2: Supplementary Fig. 16d), suggesting functional interactions between the two pathways. Interestingly, we found *IFI44L* was bound by STAT1 and STAT2 only after 6-hour treatment of IFNα, suggesting that this molecule might be a marker of sustained IFNα stimulation. DNA methylation of the *IFI44L* promoter has recently been reported to be a reliable epigenetic biomarker for SLE [35]. Similarly, *PSMB8*, which encodes a subunit of immunoproteasome, was only bound by STAT1 and STAT2 after 6-hour stimulation of IFNα or IFNγ. *PSMB8* was also found upregulated in peripheral blood cells of SLE patients (Wang et al., manuscript under review).

#### Correlation between DNA methylation and gene expression in SLE

DNA methylation is a key regulator of gene expression and we sought to find out how well they would correlate in SLE. Since there were more informative genes (significantly differentially expressed in SLE) with upregulated expression, we performed this analysis only for upregulated genes (Wang et al., manuscript under review). Before performing correlation, we assigned methylation to genes by the most significant DMR (without Bonferroni correction) within each gene promoter. We observed stronger relationship between DNA methylation and gene expression by imposing a more stringent threshold of DMR P-values for genes, as indicated by the increasing (absolute value) Pearson correlation between the reduced methylation and upregulated expression (Additional file 2: Supplementary Fig. 17a). One detailed scenario of such correlation was shown where all genes had a DMR P-value <1e-9 (Additional file 2: Supplementary Fig. 17b).

### **Supplementary Methods**

#### Analysis of overrepresentation of GWAS susceptibility loci in the DMRs and DVSs

SLE susceptibility variants with meta-analysis P-values <1e-3 and >5e-8 were selected (27,644 variants in total) based on GWAS data from UK, Hong Kong and Anhui (China) [34]. SLE suggestive risk loci were defined as regions after extension of 50kb around these variants and merging of overlapping regions. In total, 1,754 non-overlapping risk regions were obtained. Then, total number of DMRs overlapping risk loci was calculated as the observed overlapping value (O_gwas_). Then, 1000 simulations by permutation were performed to obtain a null distribution of overlaps (O_random_). For each permutation, a set of reshuffled DMR regions (i.e. same number of randomly chosen regions with the same length distribution as DMRs, excluding chromosome X to match the DMR calling procedure) were drawn from the genomic regions obtained by extending 1,000bp (the same parameter used by DMRcate to call DMRs) upstream and downstream for each CpG site interrogated from Illumina HumanMethylation450 BeadChip platform. These randomly chosen regions were overlapped with SLE risk loci. A z-score of DMR enrichment in SLE risk loci were calculated as follows:

$$Zscore=\frac{O_{gwas}-Mean(O_{random})}{SD(O_{random})}$$

The same procedure of GWAS enrichment was applied to DVSs, except that the background DVSs were randomly chosen from the 450k BeadChip (chromosome X excluded) in each permutation.

#### Analysis of overrepresentation of chromatin states in the DMRs and DVSs

The data of chromatin states [31] were downloaded from the WashU Epigenome Browser [36]. To match the cell types for the reference of genome-wide chromatin regions, we used the sample E038 for CD4+ T cell, E032 for CD19+ B cells, and E29 for CD14+ monocytes. To determine the enrichment or depletion of DMRs or DVSs in each chromatin state, we used the same process of their enrichment for the suggestive GWAS loci, with 1000 simulations for permutation test. A z-score was obtained for each chromatin state in each cell type, separately for each methylation class (hyper- or hypomethylated, and hyper- or hypovariable). Heatmaps of the z-scores were generated by R package, pheatmap, for DMRs and DVSs, respectively.

#### Analysis of DMGs bound by TFs upon interferon stimulation

To better understand the differentially enriched TFs upon various interferon stimulations, DMGs bound by those stimulated TFs (STAT1 or STAT2) were extracted from the TF enrichment analysis. The different DMGs for different treatment periods or by different types of interferons (IFNα or IFNγ) were analyzed. Three gene sets -- genes unique to either treatment and genes shared by both treatments -- were fed to ToppGene [37] for functional enrichment analysis, or GeneMANIA [38] for possible connections by protein-protein interaction, co-expression, colocalization and other functional connections.

#### Analysis of the selected DMR genes by LASSO

To investigate the relationship with clinical variables of patients, M-scores for naïve CD4+ T cells were correlated with age, ethnicity, and SLEDAI disease score. Functional Enrichment based on gene ontology and KEGG pathway were performed by clusterProfiler and ToppGene [37, 39].

#### Network analysis for the LASSO-selected DMR genes and GWAS risk genes

The GWAS risk genes were obtained from [28]. A subnetwork was extracted from the STRING database by R package igraph, containing all GWAS risk genes and their directly connected PPI neighbors. First, LASSO-selected DMR genes interacting with GWAS risk genes was defined as the overlap of DMR genes with genes within the subnetwork. Background genes interacting with the risk genes were the remaining genes in the subnetwork. Chi-squared test was used to determine the enrichment significance of interaction between DMR genes and risk genes compared with the background genes [22]. For network clustering analysis, we used Markov Clustering algorithm (MCL) to obtain PPI modules in this subnetwork. MCL was designed specifically to perform clustering based on a graph structure, such as a PPI network, and was used in many fields [40]. Distinct advantages of MCL include rapidly and accurately defining network clusters and avoiding incorrect assignments of nodes to clusters in the presence of false positive edges. The inflation parameter of the MCL algorithm was applied to control the granularity of the clustering and was set as the default value of 2.5 as in Cytoscape. After clustering, small clusters with three or fewer genes were removed. Visualized network presentation was generated by Cytoscape.

### **Supplementary Figures**

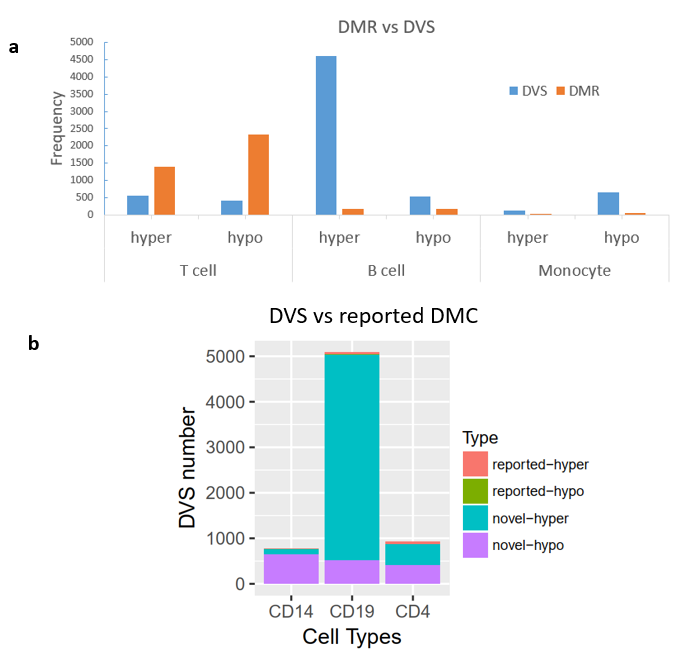

**Supplementary Fig. 1 Comparison between DVSs and DMRs or DMCs.**

(**a**) More DMRs in T cells (higher average methylation) than in B cells, but more DVSs in B cells (higher methylation variability) than in T cells. T, T cells; B, B cells; hypo, hypomethylated or hypovariable; hyper, hypermethylated or hypervariable.

(**b**) Comparison of DVSs with genome-wide significant CpGs in the original report of the same dataset.

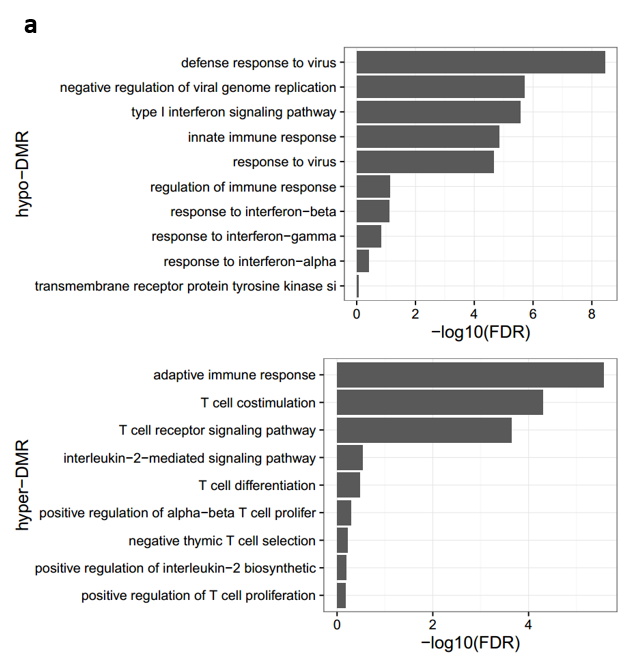

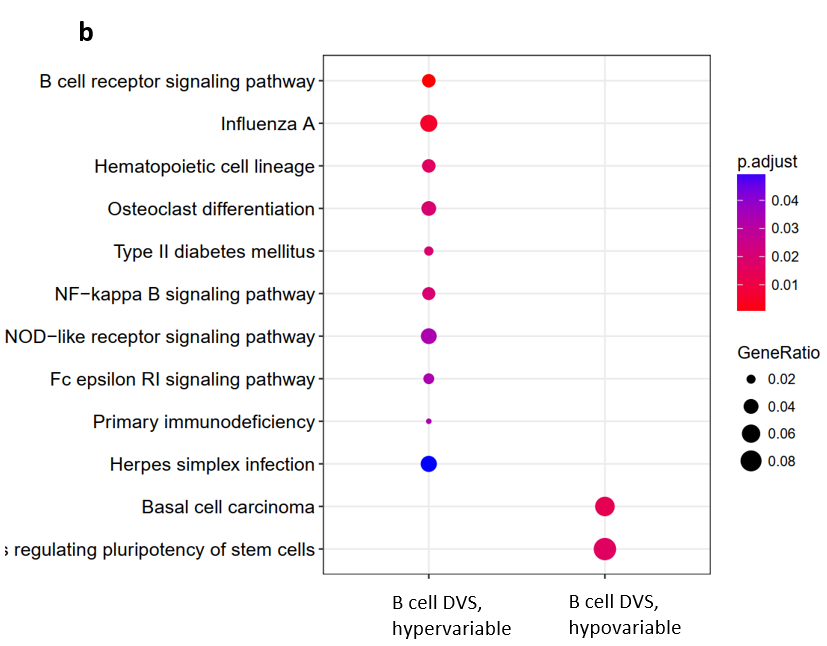

Supplementary Fig. 2 Functional enrichment for T cell DMRs and B cell DVSs.

(a) In T cells, hypomethylated genes were enriched in the innate immunity, mainly type I interferon signaling and viral defense, and hypermethylated genes were enriched in the adaptive immunity, mainly T cell receptor (TCR) signaling.

(b) In B cells, hypervariable genes were enriched in the B cell receptor (BCR) signaling.

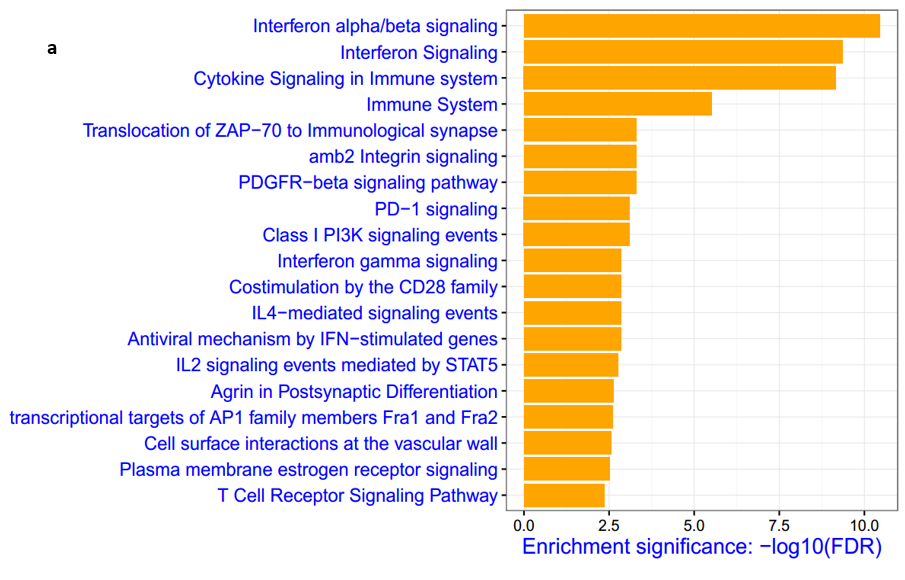

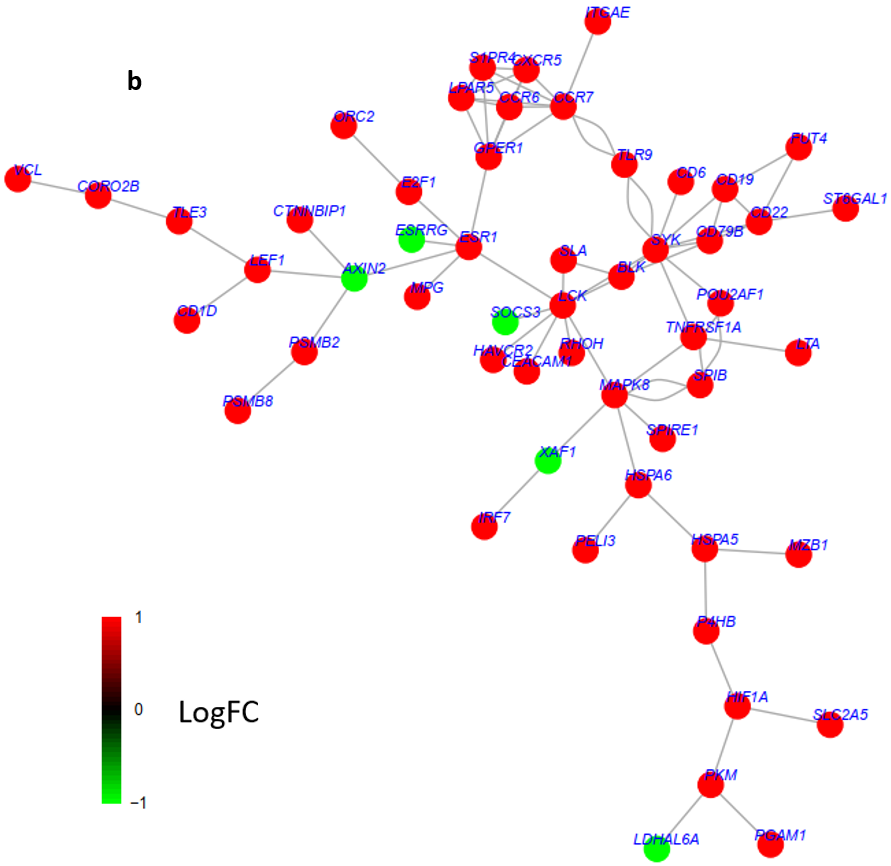

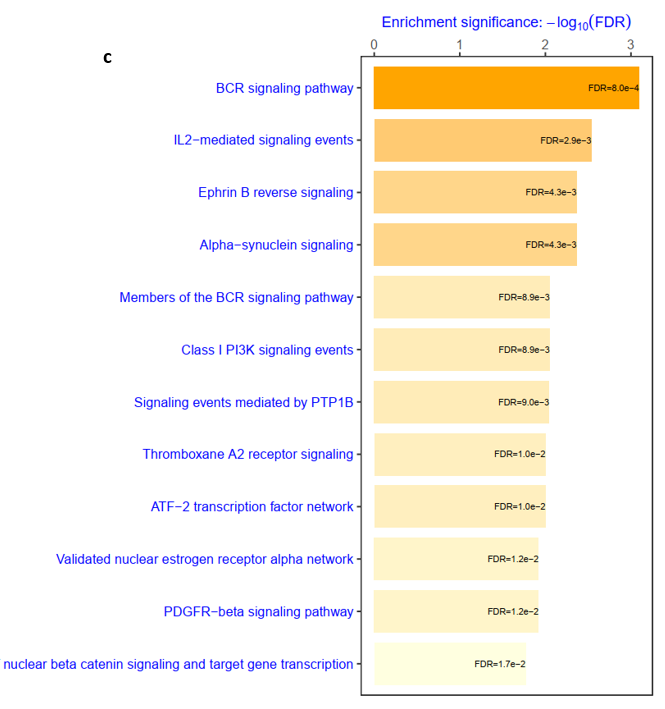

Supplementary Fig. 3 TCR and BCR signaling clusters in the PPI subnetwork.

(a) Pathway enriched for the subnetwork genes of T cell DMRs, including the PD-1 signaling pathway, ZAP-70 immunological pathway, and CD28 co-stimulation were involved in TCR co-signaling.

(b) PPI subnetwork for DVSs in B cells, where almost all genes were hypervariable (red color). Color stands for normalized log fold-change of DNA methylation in SLE compared to controls.

(c) Pathway enriched for the subnetwork genes of B cells DVSs. The BCR signaling pathway were top-ranked.

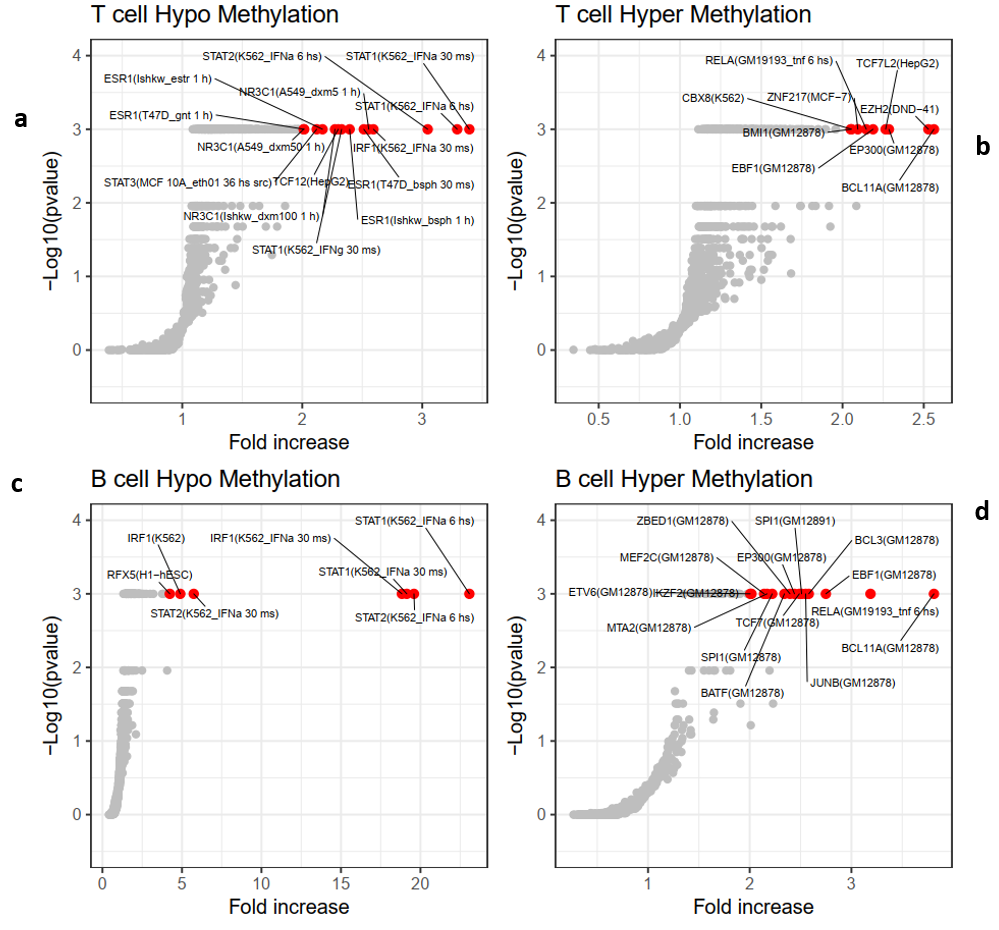

Supplementary Fig. 4 TF enrichment in DMRs. Transcription factor binding enrichment was determined by permutation test, only several top-ranked TFs were labeled for visualization.

(a, c) Mainly STAT and IRF family TFs were enriched in hypomethylated DMRs, and the fold enrichment was high in B cell hypomethylation.

(b, d) For hypermethylated DMRs, no STATs and IRFs was enriched. Instead, BATF, EBF1, BCL11A, and EP300 were top-ranked.

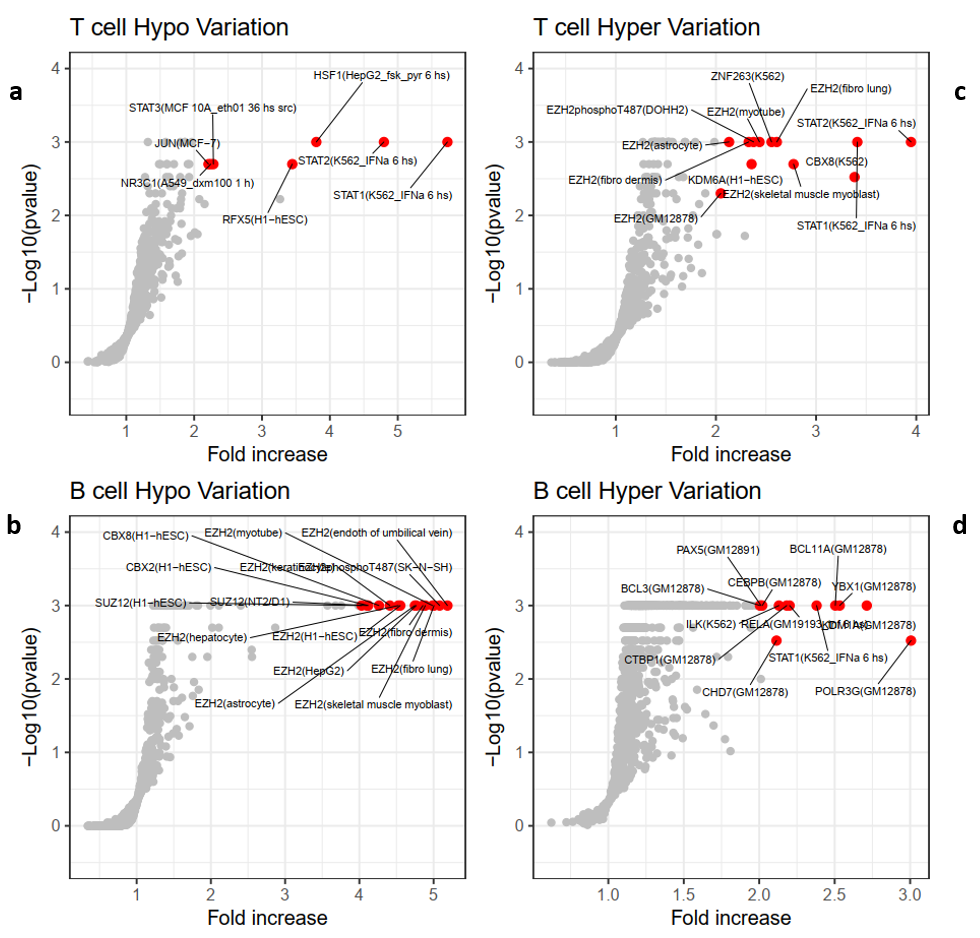

Supplementary Fig. 5 TF enrichment in DVSs. Transcription factor binding enrichment was determined by permutation test, only top-ranked TFs were labeled for the purpose of clear visualization.

(a) STAT family TFs were enriched in hypovariable DVSs in T cells.

(b, c) EZH2 was among the top-ranked TFs in hypervariable DVSs in T cells and hypovariable DVSs in B cells.

(d) Enhancer-binding TFs were top-ranked in hypervariable DVSs in B cells, including KDM1A, CEBPB, CHD7, PAX5, CTBP1, and POLR3G.

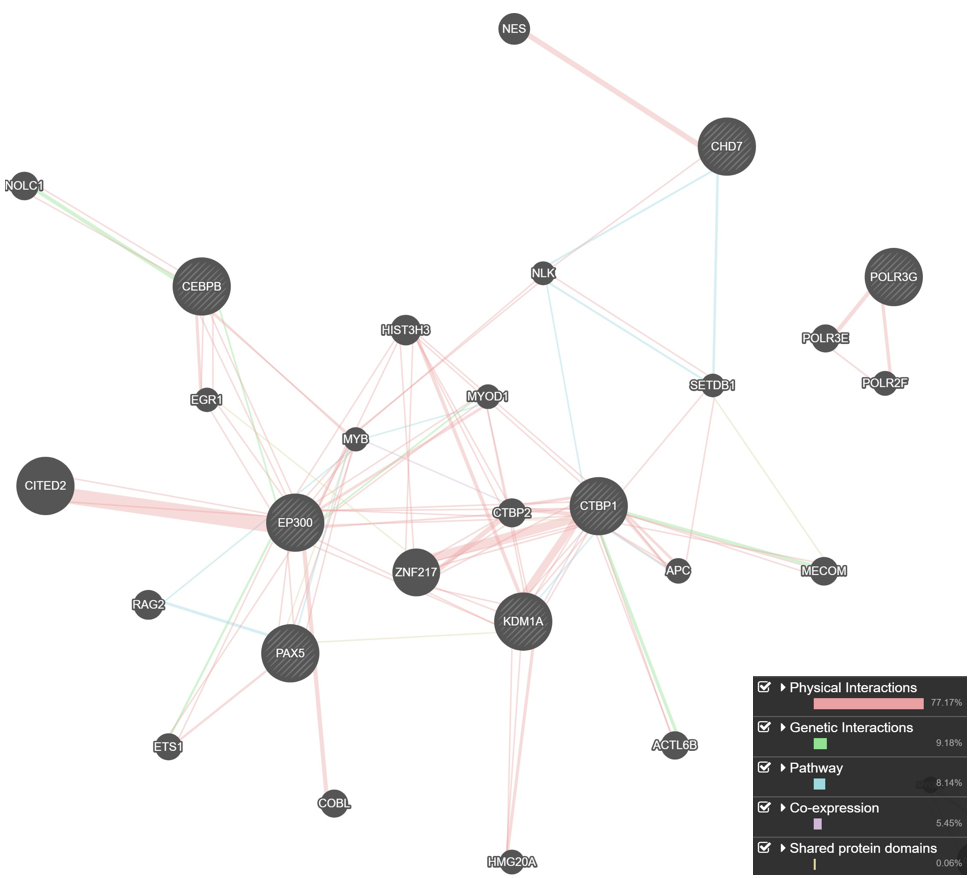

Supplementary Fig. 6 Interactions of enhancer-binding TFs enriched in hypervariable DVSs in B cells. Interactions between the enhancer-binding TFs (circles with stripes) which were enriched in hypervariable DVSs in B cells, analyzed by GeneMANIA.

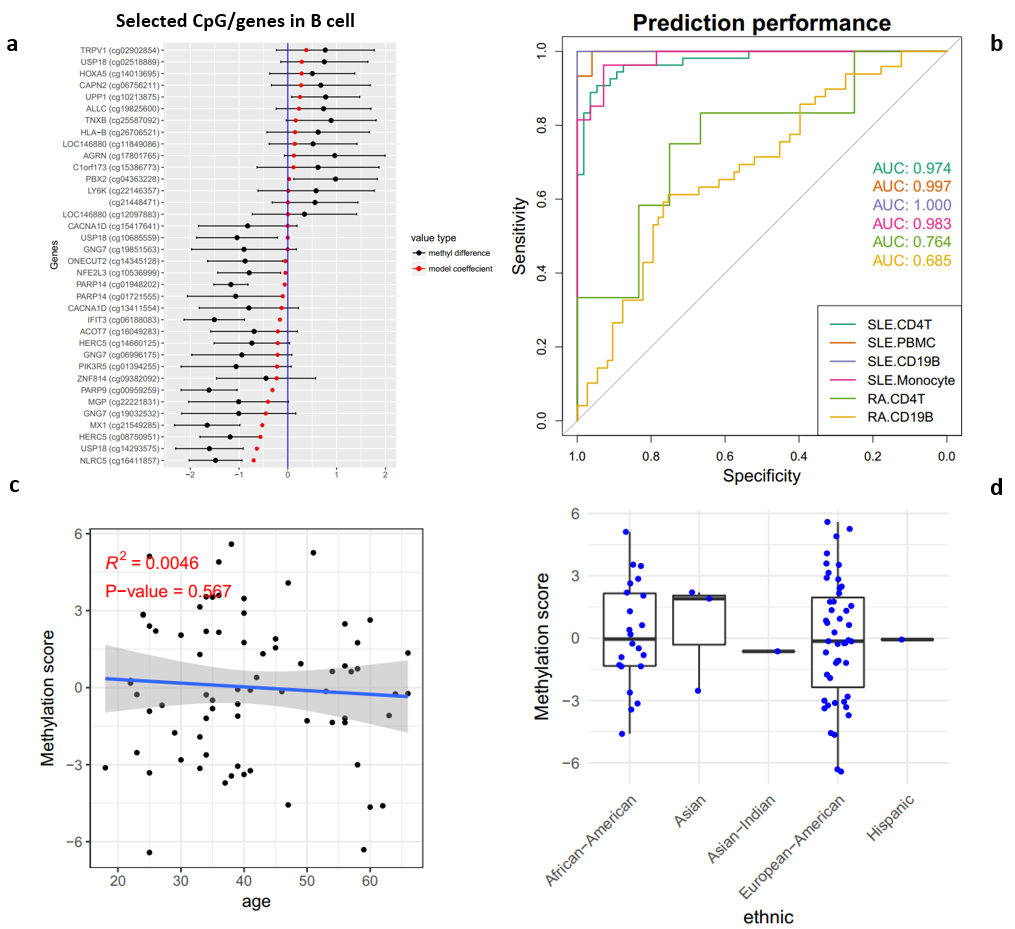

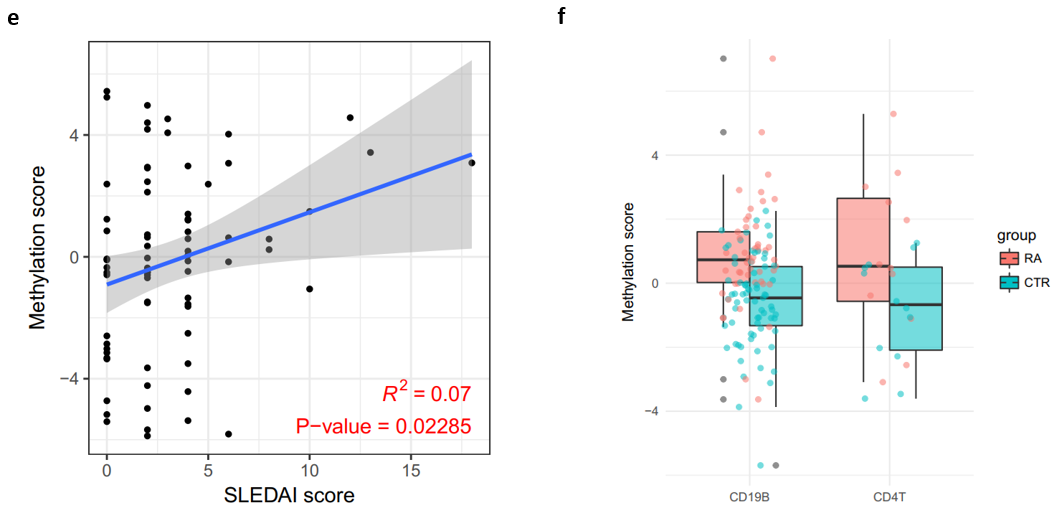

Supplementary Fig. 7 Machine learning and correlation of prediction scores with clinical variables.

(a) A list of DMR genes in B cells were selected by LASSO with different weights. Red dots indicate model coefficients and black dots with error bars shows the normalized methylation for CpGs.

(b) Model performance of (a) measured by AUC, with very good values for each cell type in SLE (0.98 on average) but much lower in RA (0.72 on average).

(c) Age was not correlated with predicted M-scores in naïve CD4+ T cells in SLE patients.

(d) M-scores showed no difference across patients with different ethnicities, using naïve CD4+ T cells.

(e) SLEDAI scores was correlated with M-scores for B cells in SLE patients.

(f) The trained model in SLE was not applicable to RA.

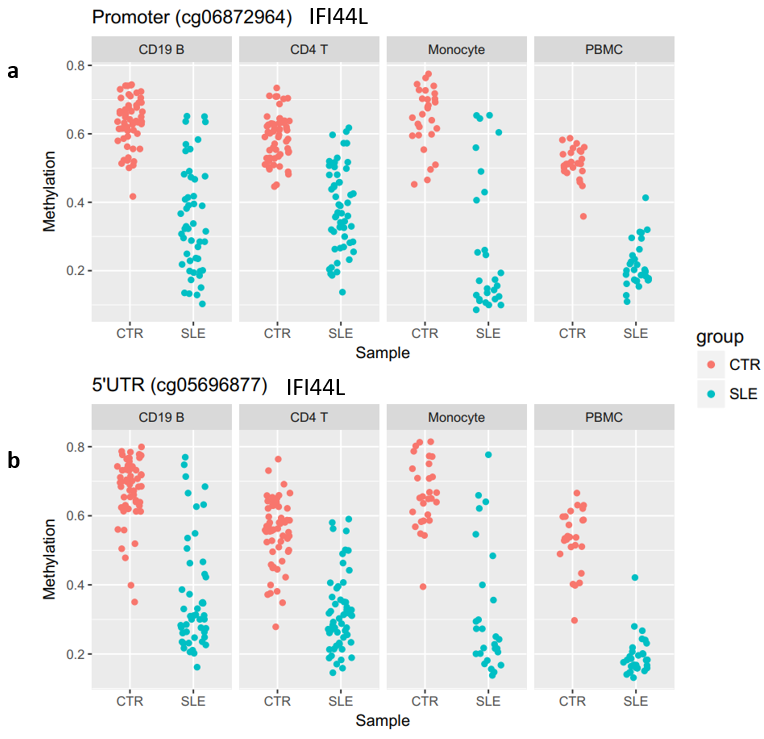

Supplementary Fig. 8 DNA methylation of two CpG sites in IFI44L in SLE and controls across different cell types. (a) cg06872964 (promoter) by Zhao et al. (b) cg05696877 (5’UTR) in this study. Only PBMC samples were clearly separable by either CpG.

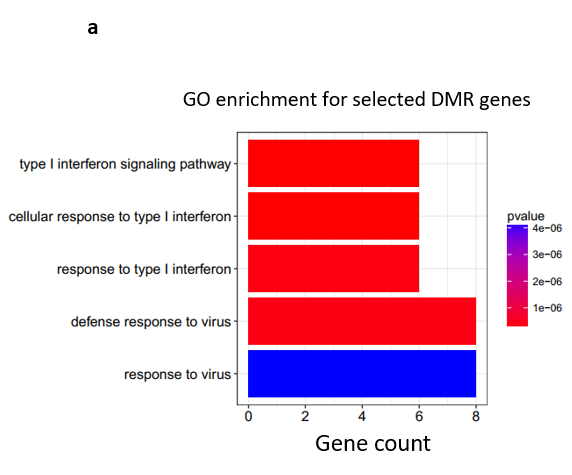

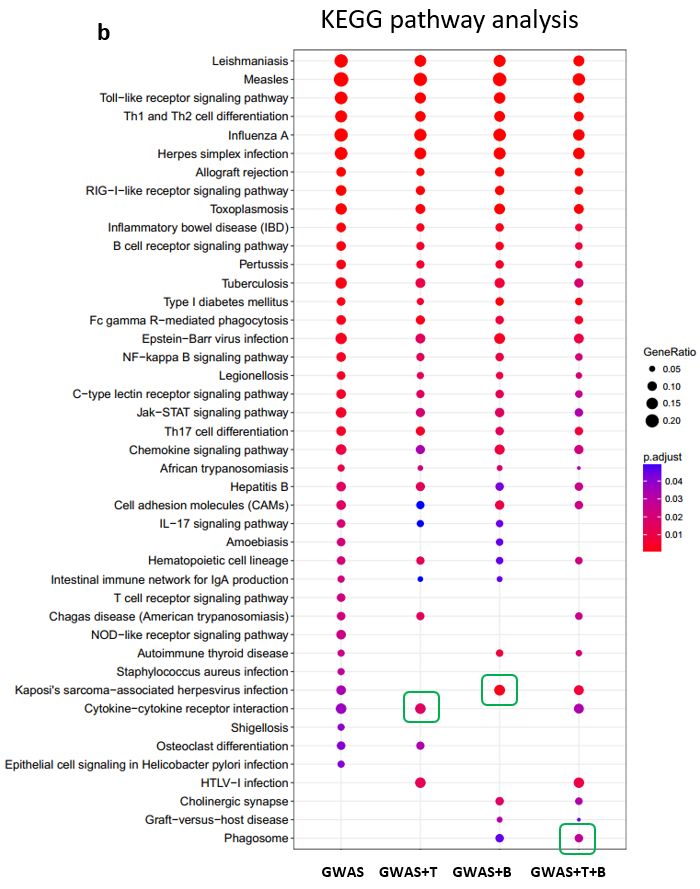

Supplementary Fig. 9 Functional enrichment analysis for DMR genes selected by LASSO.

(a) Gene ontology enrichment for selected DMR genes, which showed mainly type I interferon signalling.

(b) KEGG pathway enrichment was improved after integrating selected DMR genes into GWAS susceptibility genes, such as pathways of Phagosome, herpesvirus infection, and cytokine interaction (T: T cell DMR genes, B: B cell DMR genes).

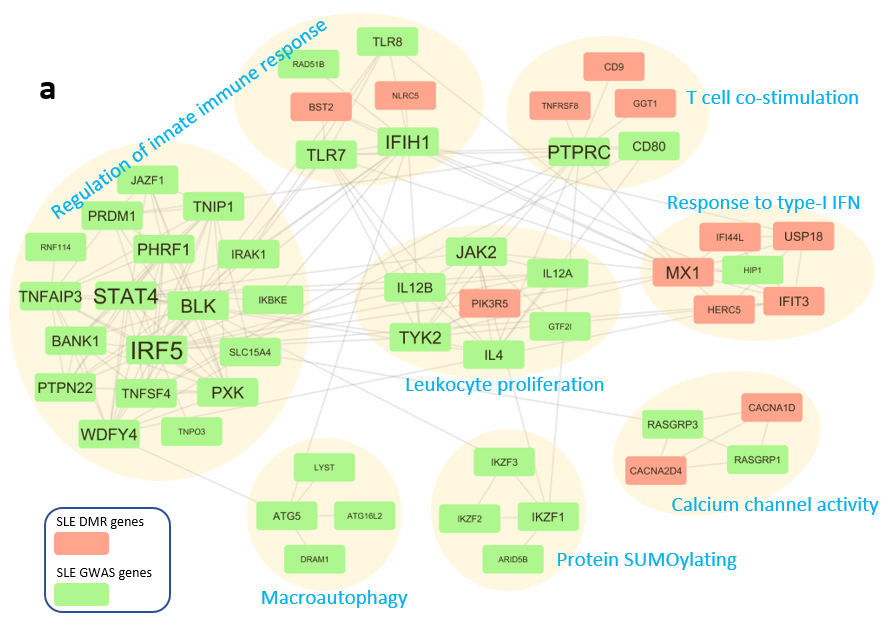

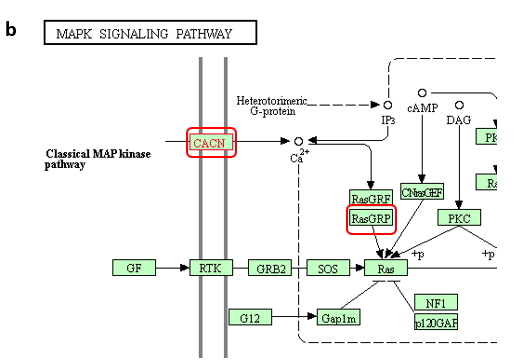

Supplementary Fig. 10 PPI network modules defined by GWAS risk genes and DMR genes selected by LASSO.

(a) PPI clusters based on a subnetwork extracted from the STRING database, containing LASSO-selected DMR genes (red box) and SLE GWAS risk genes (green box). Clusters were generated by MCL (Markov clustering) algorithm and clusters with >= 4 nodes (genes) were removed. GeneMANIA was used for pathway enrichment for each cluster.

(b) The densely-connected cluster (label: calcium channel activity) in contains two components (*CACN* and *RasGRP*) of calcium signalling inthe MAPK signalling pathway (KEGG).

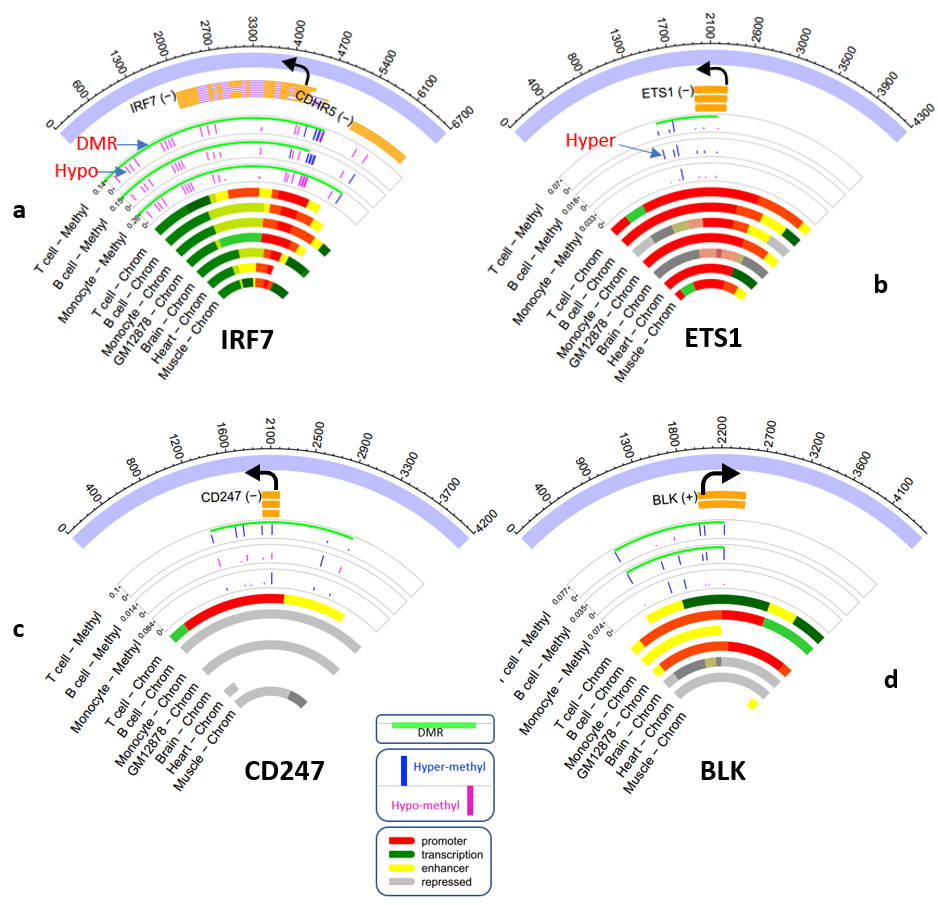

Supplementary Fig. 11 Examples of DMRs and DVSs identified in known SLE susceptibility loci.

(a-d) Four susceptibility genes were shown in detail: *IRF7, ETS1, CD247,* and *BLK*. Each had four types of genomic tracks from outmost to innermost: (1) relative genomic coordinate (base pair). (2) gene track, orange rectangles stand for exons and arrow stands for TSS. (3) methylation difference between SLE patients and controls (blue: hypermethylation, magenta: hypomethylation), with identified DMRs shown on the top of each track (green). (4) chromatin states inferred by ChromHMM.

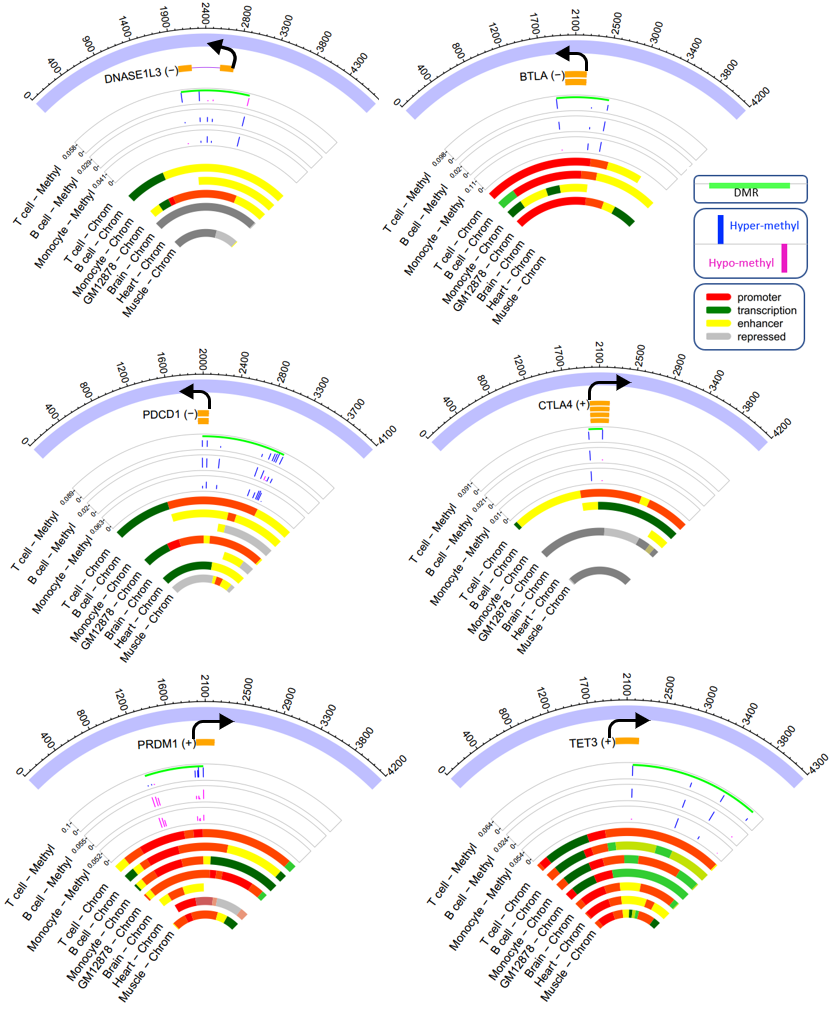

Supplementary Fig. 12 Additional representative known SLE associated genes with DMRs. Each had four types of genomic tracks from outmost to innermost: (1) relative genomic coordinate (base pair). (2) gene track, orange rectangles stand for exons and arrow stands for TSS. (3) methylation difference between SLE patients and controls (blue: hypermethylation, magenta: hypomethylation), with identified DMRs shown on the top of each track (green). (4) chromatin states inferred by ChromHMM.

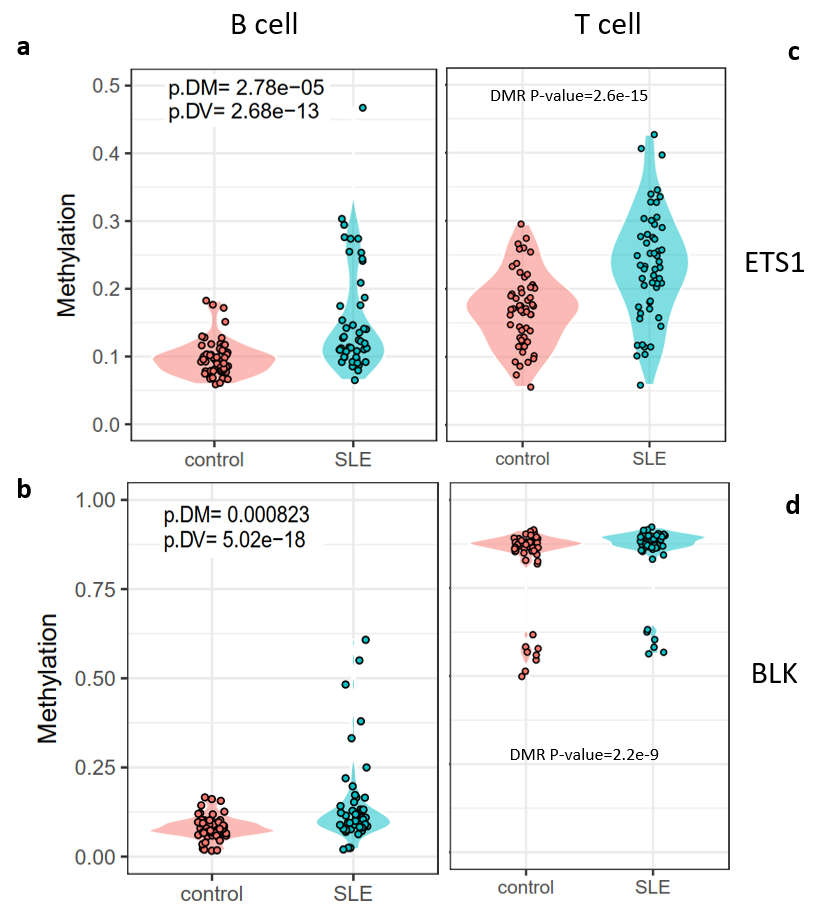

Supplementary Fig. 13 Examples of DMRs and DVSs identified in known SLE susceptibility loci.

(a-b) Two examples of susceptibility genes with B cell DVSs. *ETS1* had a differential methylation P-value of 2.78e-5 and differential variability P-value of 2.68e-13. *BLK* had differential methylation and variability P-values of 8.23e-4 and 5.02e-18, respectively.

(c-d) Dotplots corresponding to (a-b) for *ETS1* (DMR P-value 2.6e-15) and *BLK* (DMR P-values 2.2e-9) in T cells.

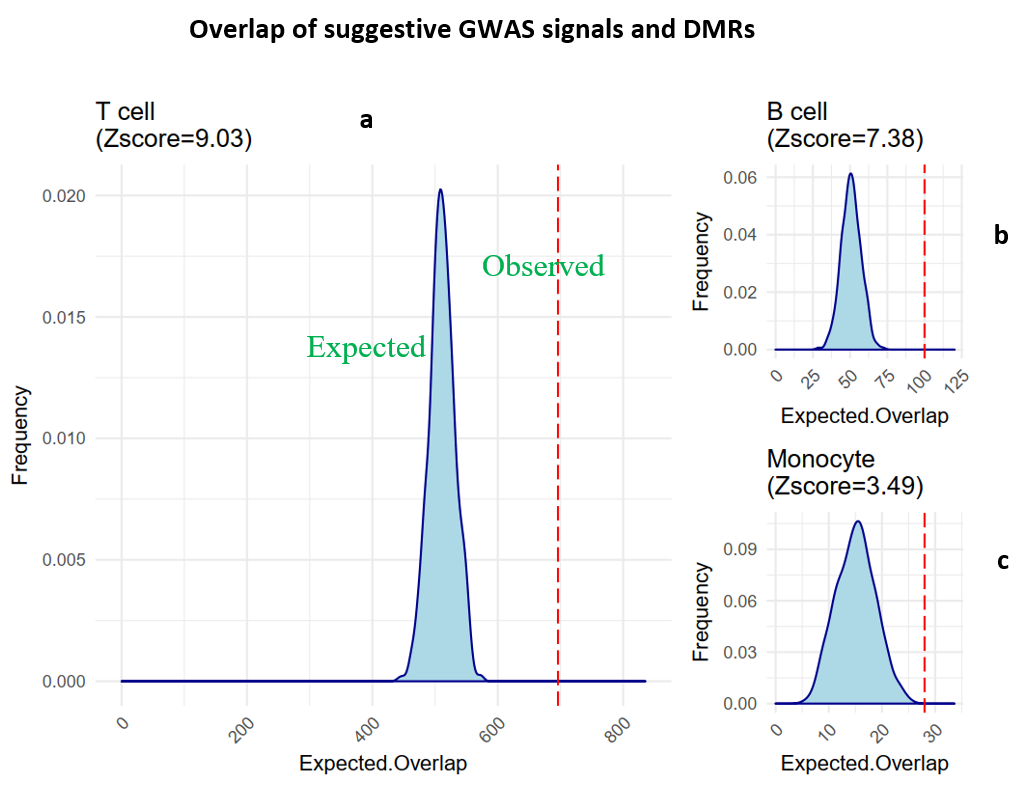

Supplementary Fig. 14 Enrichment of the suggestive GWAS loci (P-value > 5e-8 and < 1e-3) in DMRs. Enrichment was shown for T cells (a), B cells (b), and monocytes (c), as determined by permutation test. A z-score was shown for each cell type. Red dotted lines stand for observed overlap between DMRs and suggestive loci, and filled curves stand for expected overlap by random chance.

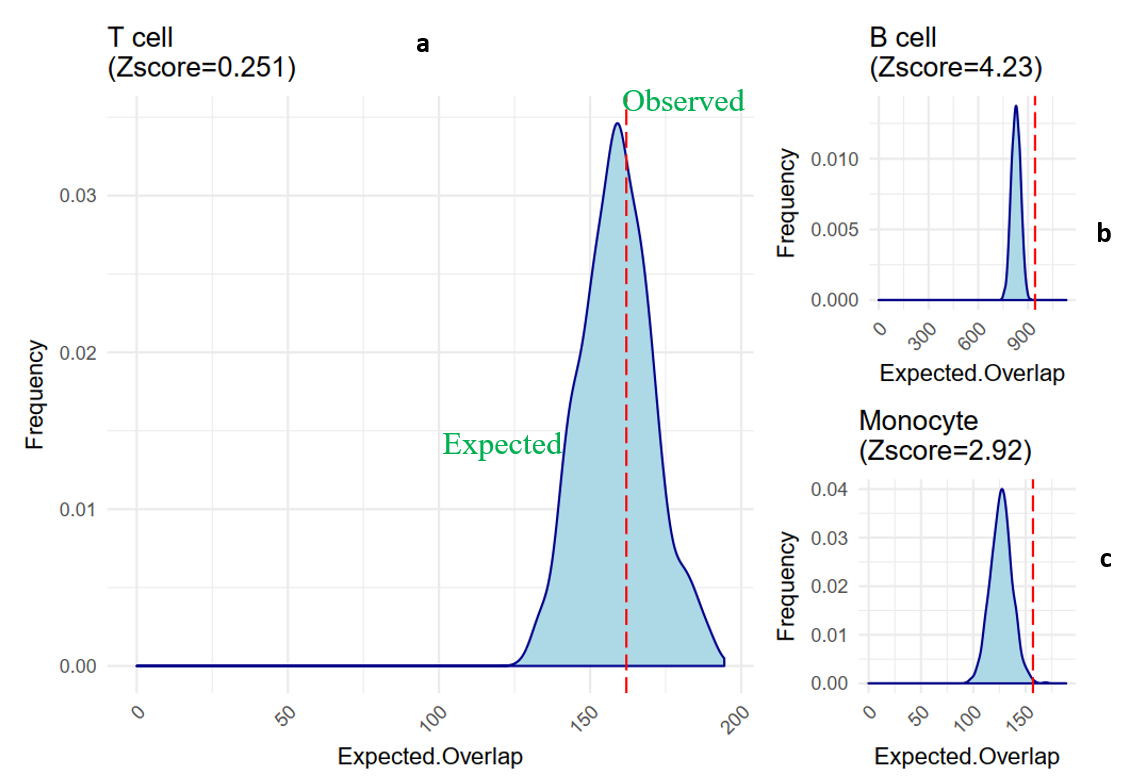

Supplementary Fig. 15 Enrichment of suggestive GWAS signals (P-value > 5e-8 and < 1e-3) in DVSs. Enrichment was shown for T cells (a), B cells (b), and monocytes (c), as determined by permutation test. A z-score was shown for each cell type. Red dotted lines stand for observed overlap between DVSs and suggestive loci, and filled curves stand for expected overlap by random chance.

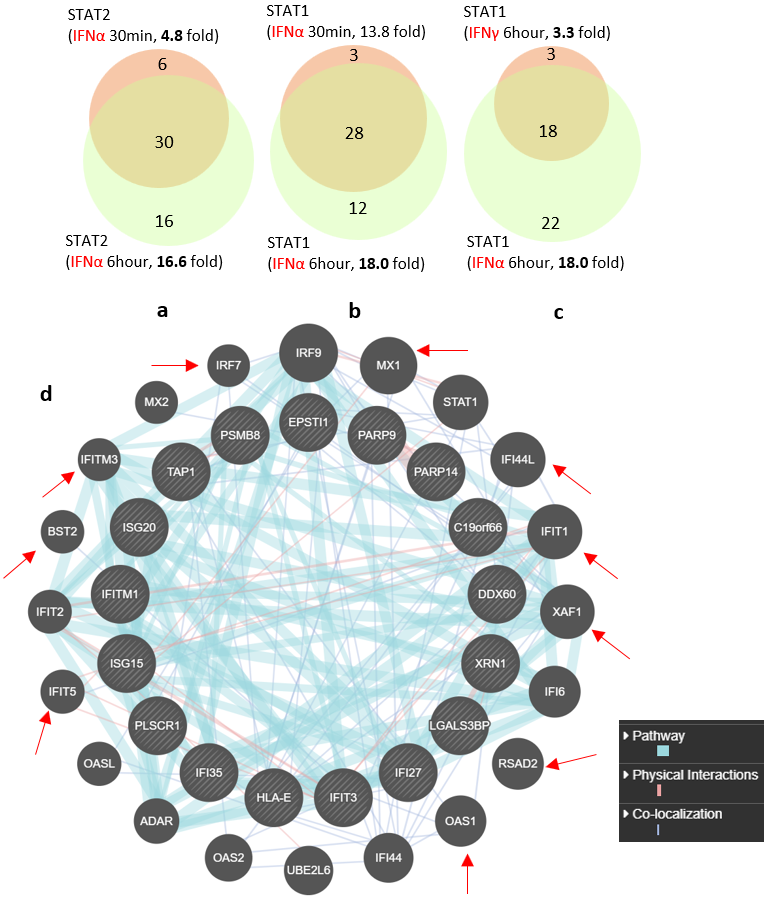

Supplementary Fig. 16 STAT1 and STAT2 upon stimulation were differentially enriched in hypomethylated DMRs in B cells.

(a-c) Venn diagrams showed the difference of bound DMGs by STAT2 upon IFNα stimulation for 30 minutes and 6 hours (a), STAT1 with IFNα stimulation for 30 minutes and 6 hours (b), and STAT1 with stimulation of IFNα and IFNγ for 6 hours (c). (d) Hypomethylated DMGs in B cells bound by STAT1 after 6-hour stimulation of IFN-gamma (inner circle) are connected with those bound by STAT1 after 6-hour stimulation of IFN-alpha (outer circle) through pathway, physical interaction or colocalization, analyzed by GeneMANIA.

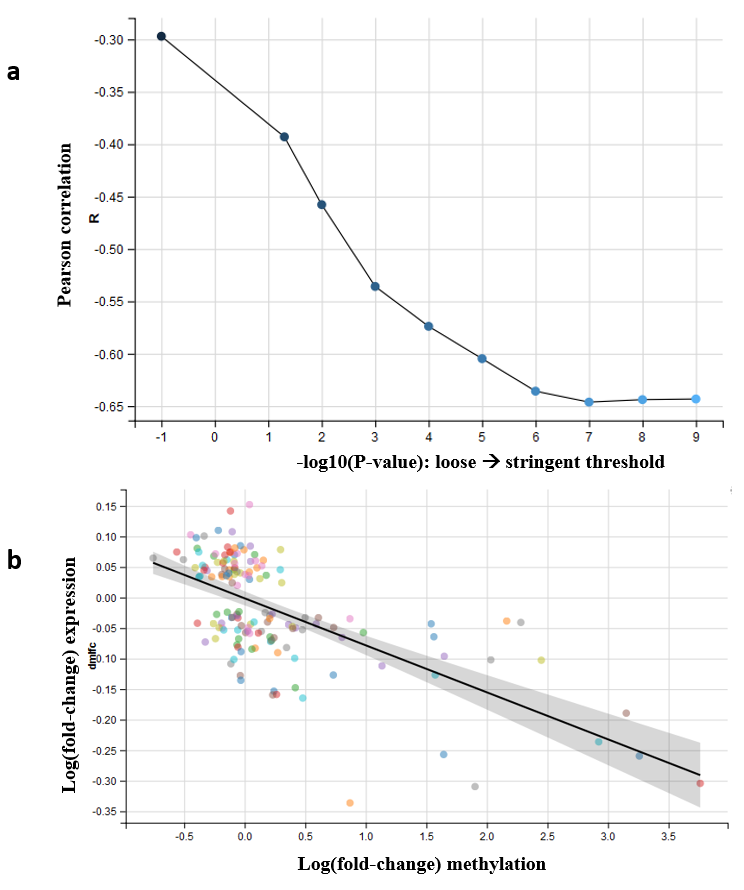

Supplementary Fig. 17 Correlation between DNA methylation and gene expression in SLE.

(a) By using more stringent criteria of DMR P-value threshold (x-axis), the negative correlation of gene expression and DNA methylation became stronger (y-axis).

(b) Under a P-value threshold of 1e-9 for DMRs, tight correlation of methylation and gene expression is shown (Pearson correlation = -0.65).

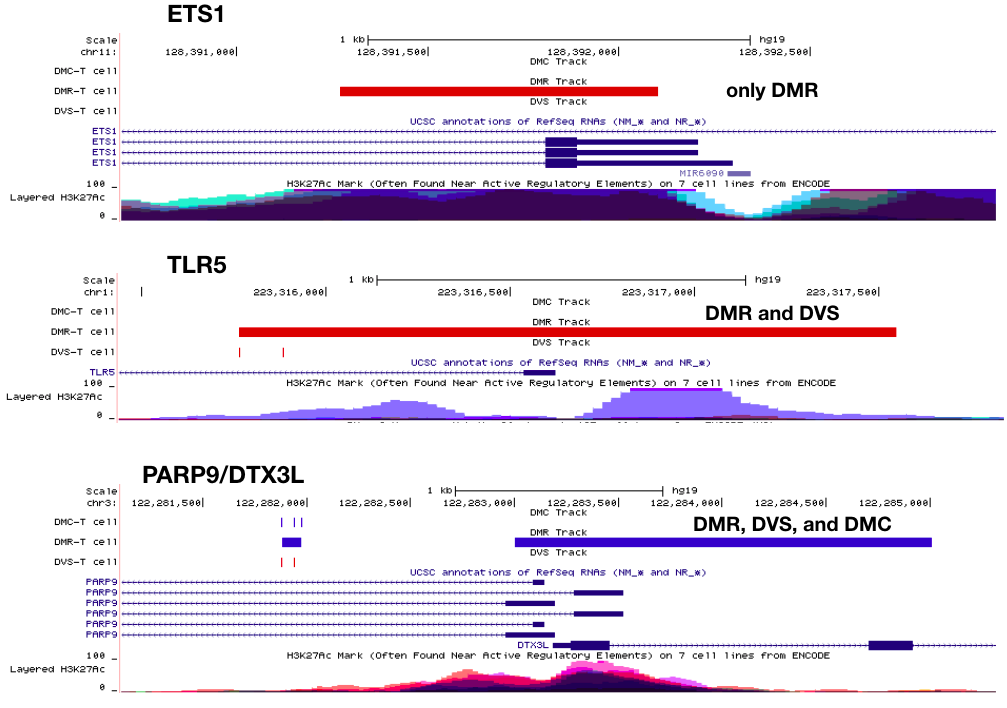

Supplementary Fig. 18 Three examples of DMR, DVS, and DMC in SLE T cells using UCSC Genome Browser.

Tracks from top to bottom are DMC, DMR, DVS, gene annotation and H3K27Ac histone marks from the ENCODE project. Only DMRs detected signal for ETS1, and both two DVSs overlapped a DMR for TLR5. Locus PARP9/DTX3L has all three types of signals. color red: hypermethylation or hyper-variability; blue: hypomethylation or hypo-variability.
